# SWAN Identification of Common Aneuploidy-Based Oncogenic Drivers

**DOI:** 10.1101/2021.05.05.442642

**Authors:** Robert R. Bowers, Christian M. Jones, Edwin A. Paz, John K. Barrows, Kent E. Armeson, David T. Long, Joe R. Delaney

## Abstract

Haploinsufficiency drives Darwinian evolution. Siblings, while alike in many aspects, differ due to monoallelic differences inherited from each parent. In cancer, solid tumors exhibit aneuploid genetics resulting in hundreds to thousands of monoallelic gene-level copy-number alterations (CNAs) in each tumor. Aneuploidy patterns are heterogeneous, posing a challenge to identify drivers in this high-noise genetic environment. Here, we developed Shifted Weighted Annotation Network (SWAN) analysis to assess biology impacted by cumulative monoallelic changes. SWAN enables an integrated pathway-network analysis of CNAs, RNA expression, and mutations via a simple web platform. SWAN is optimized to best prioritize known and novel tumor suppressors and oncogenes, thereby identifying drivers and potential druggable vulnerabilities within cancer CNAs. Protein homeostasis, phospholipid dephosphorylation, and ion transport pathways are commonly suppressed. An atlas of CNA pathways altered in each cancer type is released. These CNA network shifts highlight new, attractive targets to exploit in solid tumors.

**Highlights:**

- Copy-number alteration pathways define solid tumor biology
- SWAN is released as an integrative point-and-click pathway analysis tool
- Moderate impact drivers highlighted by SWAN validated *in vitro*
- Copy-number altered pathways associate with mutations and survival

## Introduction

Efforts to establish personalized medicine in cancer therapies have led to curative success in specific cancer types (Gambacorti-Passerini, 2008). However, progress in most lethal cancer types has been limited by the paucity of eligible patients available for testing in clinical trials. The National Cancer Institute - Molecular Analysis for Therapy Choice (NCI-MATCH) group estimated ∼9% of all cancer patients may be administered therapy based on mutation or amplification targeted data, although this already low inclusion rate does not estimate patient benefit (Flaherty et al., 2020). We previously demonstrated autophagy-loss copy-number alterations (CNAs) are druggable in high-grade serous ovarian cancer (OV). While 85-99% of OV primary tumors have a mutation in p53, there are few other canonical tumor suppressor or oncogene mutations, and none reached >10% of patients (Cancer Genome Atlas Research, 2011). It remains possible that extremely rare mutations may drive tumors like OV (Kumar et al., 2020b). The treatment feasibility issue with such exceedingly rare driver mutations is well-known: with so few patients worldwide, how can drugs be reasonably developed and approved for patient care?

CNAs are another driving factor. One seminal study in the early –omics era for cancer research showed that tumor suppressor mutations were enriched on deletion CNAs while oncogene mutations were enriched on gain or amplification CNAs (Davoli et al., 2013). We similarly found that networks built from molecular pathways and scored by CNA data were suppressed, with known tumor suppressors as network hubs, or elevated, with established oncogenes as network hubs (Delaney et al., 2017). Within the most suppressed OV pathway network, the autophagy pathway, we identified *BECN1* and *LC3B* as the most influential gene hubs. Suppression of either autophagy gene sensitized cells to autophagy inhibitors chloroquine phosphate or nelfinavir mesylate. In a platinum resistant patient-derived xenograft (PDX) model, we found autophagy targeted drugs could completely abolish observable tumor burden, even when dual platinum-taxane combination therapy had no effect on the PDX model. These results demonstrate tumor CNAs, even monoallelic CNAs, are pharmacologically targetable. Pharmacologic treatment of CNAs may be amenable to removal of early pre-cancerous cells for some tumor types. For example, clear cell renal cell cancers often lose chromosome 3p years prior to development of disease (Mitchell et al., 2018). CNA losses persist during disease progression, suggesting that they remain as biological drivers or at least persistent vulnerabilities (Mamlouk et al., 2017; Patch et al., 2015; Wedge et al., 2018).

CNAs rarely encompass a single gene. Entire chromosomes are often altered within solid tumors, creating CNAs across hundreds of genes with a single genetic alteration. Few studies have adequately addressed this background noise problem of thousands of gene-level CNAs in each tumor, many of which are passengers, for both logistical and conceptual reasons. It is arduous or infeasible to model aneuploid events in cellular and mouse models, precluding causal genetic studies. However, a handful of well-controlled studies have been completed. *TP53*-adjacent genes, *EIF5A* and *ALOX15B,* contribute to tumor formation and progression. In a lymphoma Eμ-Myc pre-B cell mouse model, suppression of these genes by shRNA independently increased lethality (Liu et al., 2016). Chromosome 8p is often lost in cancers while chromosome 8q is often gained. Here we adopt the definition of aneuploidy as changes encompassing entire chromosome arms (Ben-David and Amon, 2019); 8p loss or 8q gain are both independent aneuploid events. To model 8p loss in breast cancer, 8p loss was engineered in non-malignant MCF10A cells (Cai et al., 2016). While 8p deletion did not induce transformation, cells exhibited increased invasiveness and elevated mevalonate metabolism. In most cases, a single aneuploid chromosome causes a cell cycle delay. Accordingly, the 8p deletion exhibited this phenotype. Furthermore, a 3p deletion commonly found in lung cancers decreased proliferation in an immortalized lung epithelium AALE cell line, although cells eventually adapted (Taylor et al., 2018). Aneuploidy itself leads to increased usage of the proteasome and autophagosome machinery (Santaguida et al., 2015), indicating metabolic inefficiency. Transcriptomic compensation for aneuploidy is rare (Torres et al., 2016) and protein levels correlate well with CNAs (Zhang et al., 2016). Aneuploidy can lead to slowed tumor growth in Ras mutant xenografts (Sheltzer et al., 2017). However, under specific nutrient or signaling conditions, select aneuploid events increase cellular fitness. In serum starved cells, trisomy 13 cells exhibit greater fitness than control cells (Rutledge et al., 2016). These examples directly demonstrate CNAs drive cancer in specific selective conditions.

Here we developed a new pathway network algorithm, Shifted Weighted Annotation Network (SWAN), designed to handle high-noise biological data, such as monoallelic CNAs spread across the genome. We report SWAN analysis of 10,395 tumors studied by The Cancer Genome Atlas (TCGA) from 31 cancer types and 4,925 pathways. We demonstrate SWAN prioritizes known tumor suppressors and oncogenes within CNAs. SWAN further characterized 24 high-confidence novel multi-cancer oncogenes. Molecular pathway suppression caused by loss of tumor suppressor genes were prevalent across tumor types, representing potentially targetable vulnerabilities. We show biological validation of a tumor-specific elevated pathway, peroxisome biogenesis, and a multi-cancer suppressed pathway, cadmium response. We release an online CNAlysis Atlas and easily accessible web-based SWAN pathway analysis tools.

## Results

### Oncogenic CNAs complement other mutations

Copy-number alterations function as second-hit drivers for oncogenesis, arising after canonical drivers like *KRAS* or p53 mutation. Previous studies of normal tissue found normal epithelium contains oncogenic driver single-nucleotide variants (SNVs) and insertion-deletion (indel) mutations (Lee-Six et al., 2019; Martincorena et al., 2018; Martincorena et al., 2015; Moore et al., 2020; Yoshida et al., 2020). Stromal mechanisms may initially prevent further expansion of these clones (**Figure 1A**). Secondary drivers are necessary to escape local arrest and expand the tumor microenvironment. Initial oncogenic mutations allow for the slow development of clonal CNAs over generations of random chromosome gains and losses (**Figure 1B**). In support of the second-hit CNA driver hypothesis, our analysis of TCGA tumors reveals 5-50% of solid tumors contain only oncogenic mutations found in normal tissue clones, out of 251 previously identified driver SNVs (**Figure 1C**). Solid tumors have 15-70% of each tumor genome altered by CNAs, with a median alteration of 39% of the genome (**Figure 1D**). The scale of established oncogenes (OGs) and tumor suppressor genes (TSGs) on gain or loss CNAs, respectively, range from 20 to 30 of each in solid tumors (**Figure 1E**). CNAs of known driver genes are a hallmark of solid tumor genetics.

**Figure 1.**
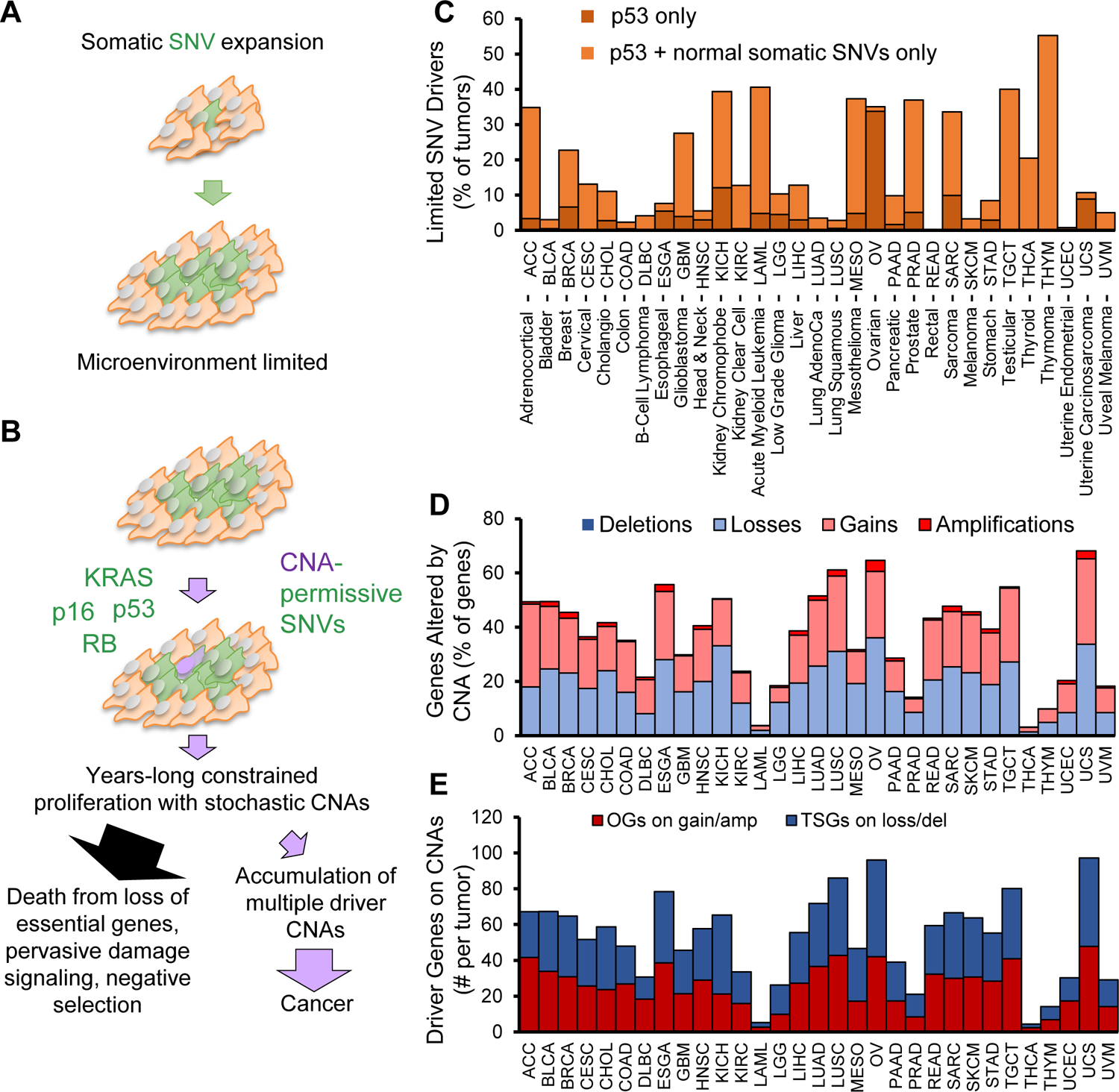
Copy-number alteration drivers are present in tumors with insufficient SNV drivers. (**A**) Model for limited growth of epithelial cells with normal-tissue clonal SNVs. (**B**) Model for oncogene accumulation by CNAs to drive cancer initiation and progression. (**C**) TCGA tumors were analyzed for the number of tumors with insufficient SNV drivers. Tumors were queried for 251 Tier 1 COSMIC oncogenes and tumor suppressor gene mutations. Tumors with only p53, or with p53 and mutations commonly found in normal human epithelium (includes *NOTCH1-3*, *FGFR3*, and *FAT1*) and no other COSMIC OG or TSG mutation are plotted as a percent of all tumors queried. (**D**) Frequency of CNAs in the same cancer types. (**E**) COSMIC Tier1 cancer genes overlapping deletion CNAs (for TSGs) or amplification CNAs (for OGs).

### Design of Shifted Weighted Annotation Network (SWAN) analysis

Biological pathways contain multiple genes that are typically located on multiple chromosome arms. Thus, tumors with different chromosome content may nonetheless upregulate the same pathway if genes within the same pathway are altered in the same direction (either losses or gains). These individual genes may differ between patients, yet the pathway is nonetheless similarly altered in flux. To quantify and prioritize such complex changes in biological data, we developed SWAN analysis.

SWAN analysis is broadly applicable to any gene-level data set and is similar conceptually to Gene Set Enrichment Analysis (GSEA), but includes the addition of interaction and phenotypic data to improve the testing of suppression or activation pathway hypotheses by forming weighted pathway networks. In SWAN, pathway-specific elevation and suppression hypotheses are independently tested and compared to a randomized null hypothesis. Permutations of gene-level data are done for each tumor (1,000 random pairs in this study) to generate null distributions specifically relevant to each sample and pathway (**Figure 2A**). Control sample data may additionally be utilized. SWAN performs statistics on experimental sample network shifts relative to permuted or sample controls, using a single network shift score per sample for large datasets (N ≥ 15), or using individual gene shifts per tumor for smaller sample sizes. Significance thus scales with sample size, not with the number of permutations. SWAN performs additional steps to reduce noise in the results (**Figure S1** and Methods). The precise details of SWAN calculations can be found in Methods, the supplied code, tutorial videos, and the documentation provided with the program. For this study, the “annotation” is genes, but investigators may use SWAN in the context of lncRNAs, miRNAs, replication origins, or any other annotation of the genome, provided that the investigator can supply pathway lists and optional interaction data.

**Figure 2.**
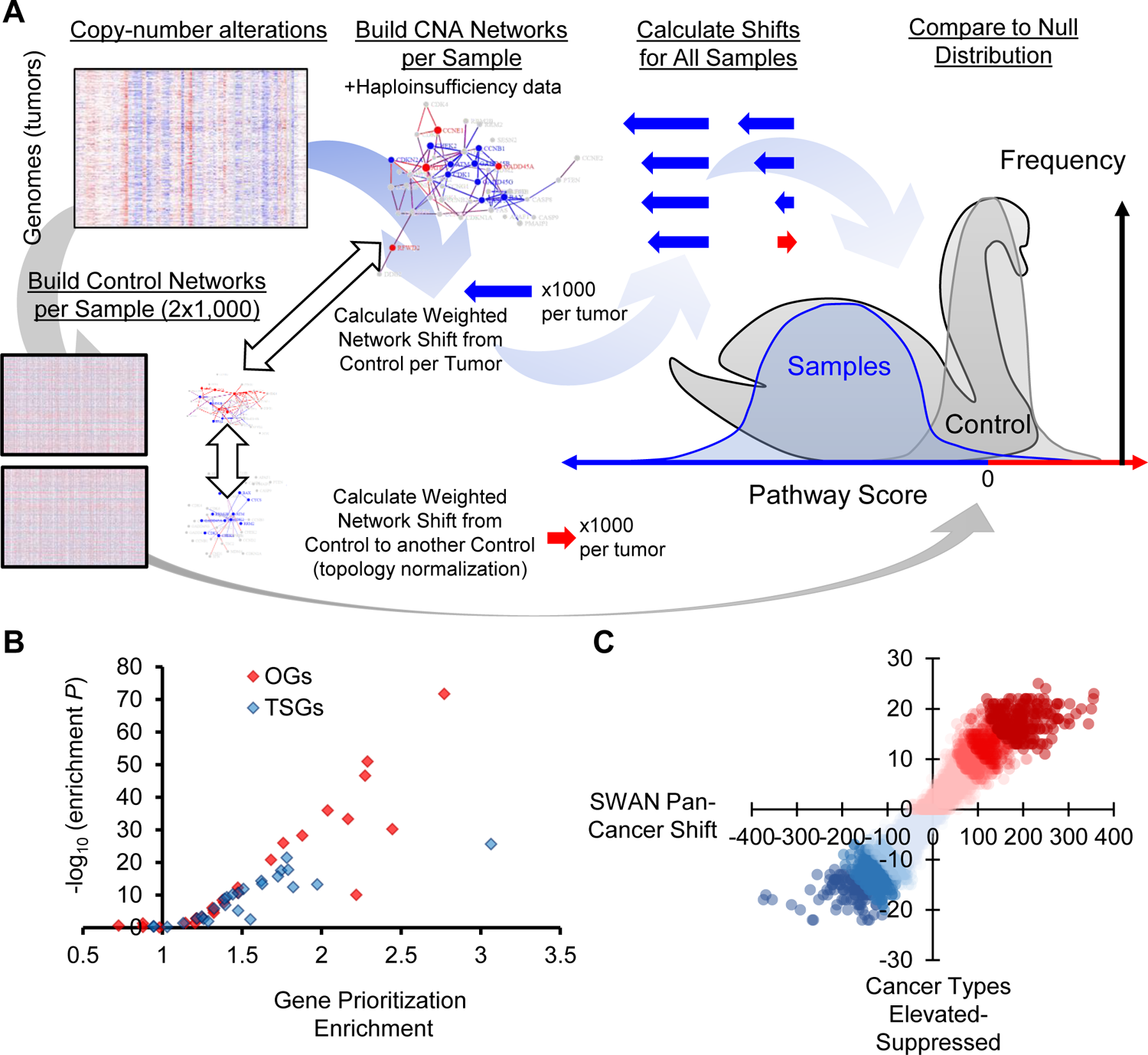
Design of Shifted Weighted Annotation Network (SWAN) pathway analysis tool and pan-cancer results. (**A**) A conceptual diagram of SWAN calculations. Raw data in this pan-cancer analysis is CNAs. Pathway networks are then built utilizing protein-protein interaction data and haploinsufficiency data. Sample network scores are compared to paired-shuffled control data. Details are found in SWAN documentation and Methods. (**B**) Plot of the statistical enrichment of known OGs and TSGs on elevated or suppressed pathways in each of the 26 QC-compatible tumor types studied. (**C**) Plot of the number of pathways which are identified among 31 cancer types as elevated or suppressed relative to the sum of SWAN shifts.

### Quality control of SWAN for cancer data

CNA analyses have previously focused on segments of DNA that are significantly altered in tumors. Extreme amplifications of genes like *MYC* or *EGFR* and homozygous deletions of *CDKN2A* are highlighted in previous studies due to the high-noise nature of aneuploidy patterns (Beroukhim et al., 2010; Smith and Sheltzer, 2018; Zack et al., 2013). These analyses often ignored the biological changes caused by the 90% of tumor CNAs: removal or duplication of a single allele. Using GSEA, one of the most popular pathway analysis tools with >10,000 citations (Subramanian et al., 2005), OV had only a single significantly elevated Kyoto Encyclopedia of Genes and Genomes (KEGG) pathway, “complement and coagulation cascades”. Applied to 26 cancer types, very few, if any, pathway alterations were found by GSEA across all KEGG pathways (**Table S1, Figure S2A**). This is unlikely to be the real biological situation for tumors containing clear patterns across chromosome arms. SWAN was designed to circumvent this false-negative issue using phenotype-layered networks.

To assess phenotype importance in appropriately defining tumor CNA genetics, we performed multiple pan-cancer SWAN calculations across 26 quality control (QC) compatible cancer types studied by TCGA with 4,925 pathways (KEGG, Gene Ontology [GO], Hallmark, and Reactome). As a positive control, we used TSGs and OGs from COSMIC’s Tier 1 Cancer Gene Census (Sondka et al., 2018). SWAN identifies the most influential suppressed genes within suppressed pathways and the most influential enhanced genes within elevated pathways. If working appropriately in a cancer context, suppressed pathways should have TSGs prioritized and elevated pathways should have OGs prioritized. There was a significant (*P* ≤ 0.05) enrichment of TSG prioritization within suppressed pathways in 23 of 26 tumor types and a significant enrichment of OG prioritization amongst elevated pathways in 22 of 26 tumor types (**Figure 2B**). Cancer types without significant enrichment had unusually low CNAs, consistent with the hypothesis that CNAs are not strong drivers amongst all tumors in these cancer types. Applying identical quality control to GSEA and SWAN KEGG pathway analysis, SWAN outperformed GSEA in prioritizing driver genes in 22 of 26 tumor types (**Figure S2B**). Amongst all 31 cancers studied, no pathway was altered in one direction in every tumor type and there was a wide distribution of pan-cancer pathway shifts to single-cancer shifts (**Figure 2C**).

To test if creation of phenotype networks aided in the prioritization of TSGs and OGs, three additional pan-cancer SWAN analyses were performed. First, removal of haploinsufficiency and triploproficiency scoring yielded a moderate and consistent decrease in SWAN’s ability to prioritize TSGs and OGs across tumors (**Figures S2C, S3**). Second, the protein-protein interactions (PPIs) used to build pathway networks were removed. Removal of PPIs substantially reduced the ability of SWAN to correctly prioritize TSGs and OGs. Third, removal of both phenotypes from SWAN completely abrogated prioritization of TSGs and reduced prioritization of OGs by 87%. These data support the use of phenotype information, particularly haploinsufficiency and PPIs, to aid in the analysis of CNAs in cancer. Noting that COSMIC-annotated OGs were better prioritized in SWAN than TSGs, we postulate that true TSGs on CNA regions may be “moderate or low impact” TSGs in that multiple TSG deletions are necessary for a stronger pro-proliferative effect (Davoli et al., 2013; Liu et al., 2016). To test this, QC was also performed using STOP and GO genes as annotated from at least two sources (see materials). Unexpectedly, STOP genes were not more enriched than known TSGs, potentially due to a lack of tissue-specific information ((Sack et al., 2018), reference Figure S2C). Taken together, these QC tests show SWAN appropriately prioritizes genes most likely to act as true TSGs or OGs within CNA data across cancers.

### Development of point-and-click integrated network analysis web platform

SWAN was designed to be useful to statisticians, bioinformaticians, and molecular biologists alike. Specifically, SWAN is available in two forms: as an R package with standalone code (for statisticians and bioinformaticians, at https://github.com/jrdelaney/SWAN) and as a hosted website (for everyone, https://www.delaneyapps.com/#SWAN). R standalone code is streamlined for minimal memory use with fast computation time. The point-and-click applications are optimized for minimal user input with logical defaults and downloadable example input files. To enable use from non-programmers, all input data are designed to be simple tab or comma delimited spreadsheets readily manipulated in Excel or Google Sheets. Segment to gene mapping, mm9 to mm10 or hg19 to hg38 conversion, and basic −2 to 2 scaling and normalization capabilities were built into the SWAN Data Groomer. The CNAlysis Atlas is pre-loaded with TCGA data to enable users to query their pathway(s) of interest without any time investment in setting up software. This includes SWAN analysis of 10,395 tumors from 31 cancer types. Shiny App versions of the statistical SWAN software were developed to enable molecular biologists with no programming experience to readily perform these advanced SWAN network analyses, including integrated RNA and mutation analyses. As such, a priority on graphical and intuitive outputs was made. Entry-level readme files and instructional videos are linked within the Shiny App.

SWAN was developed as a generalizable tool, not just for cancer. Any log_2_ normalized data or copy-number data can be used. Mouse and human pathways are included by default. Any organism may be queried using user-uploaded files. Other uses of SWAN include weighting by mutations, RNA expression, or protein modification rates, among many other possible weighting criteria for the networks. Custom interactions other than protein-protein interactions from BioGRID, such as genetic interaction data, can be uploaded and used to generate networks. Interaction-independent analyses can be performed if desired.

To demonstrate broad utility, we provide an example of SWAN use in a generic molecular biology context. We first used SWAN to analyze RNA-seq data from control or JQ1-treated human primary myofibroblasts (Suzuki et al., 2020). JQ1 inhibits the DNA localization of the BET family of proteins, traditionally identified as major epigenetic regulators. BET inhibition has become a strategy for tumor treatment; however, the fundamental biology of BET proteins (BRD2/3/4) is still being unraveled. By querying Reactome pathways in SWAN, we identified similar conclusions as the original authors: both collagen formation and extracellular matrix organization pathways were significantly down regulated in JQ1-treated cells (**Figure S4A**). We noted several other related suppressed pathways, including DNA replication, activation of the pre-replicative complex, and unwinding of the DNA. To further investigate these results, we next analyzed RNA-seq data from three other publications using multiple forms of BET inhibition across several cell lines (Garcia-Carpizo et al., 2018; Nagarajan et al., 2017; Ren et al., 2018). In all these independent datasets, BET inhibition suppressed expression networks of genes involved in the activation of the pre-replicative complex (**Figure S4B**). These results were recapitulated in a genetic knockdown of *BRD4* (**Figure S4C**). BET inhibition has been suggested to affect genes involved in DNA replication (Du et al., 2018), however, there is little literature detailing these findings. Interestingly, several proteins involved in the pre-replication complex, including CDC6, MCM5 and MCM7, have been implicated as BRD4-interating partners (Zhang et al., 2018). Indeed, BET inhibition has resulted in DNA replication stress (Bowry et al., 2018) and replication re-initiation (Zhang et al., 2018). BET proteins are also implicated in the regulation of proliferating cell nuclear antigen unloading (Kang et al., 2019). The SWAN analysis presented here underscores the importance of BET proteins in the biology of DNA replication and serves as an example of SWAN usage outside of cancer genetics.

### Integrative analysis

While the current study is primarily intended to provide new light into tumor genetics using solely CNA data, there is community interest in allowing for integrative analyses. SWAN enables the routine integration of RNA and mutation data, amongst other types of layered data (**Figure S2D**). By comparing SWAN shifts using CNA data alone to those shifts produced by RNA layered onto CNA data, outlier pathways with exceptional RNA modulation can be identified. In bladder cancer, we found an upregulation of xenobiotics and drug metabolism RNAs and fatty acid degradation relative to DNA copy number (**Figure S2E**, red), whereas the spliceosome pathway had a reduction in RNA relative to DNA copy number (**Figure S2E**, blue). Mutation shifts were less striking due to the infrequently consistent mutation events for driver genes in most cancer types. *TP53* is a rare exception in that it is commonly mutated in entire cohorts, shifting the p53 signaling pathway away from the null in an otherwise well-correlated pan-pathway analysis (**Figure S2F**). Overall transcription shift correlations across 4,912 pathways were high when RNA was scored only if in the same direction as CNAs, while specific cancer types had widely different RNA shifts when scored additively with CNA data (**Figure S5**).

### Identification of known and novel oncogenic pathways

In the SWAN pan-cancer CNA analysis, the hyperosmotic response was the most commonly elevated pathway (25 of 31 cancers elevated) (**Figure 3A, Table S2**). The most common SWAN impactful genes within the hyperosmotic response included *ARHGEF2*, *AQP1*, and *RAC1*. *RAC1* is a tier 1 COSMIC oncogene and ARHGEF2 is required for RAS-mediated oncogenesis. AQP1 is best known for its role in enabling water transporting along an osmotic gradient in kidney proximal tubules, but is also implicated in endothelial cell migration (Verkman, 2011). The second most commonly elevated pathway was epidermal growth factor receptor signaling (24 of 31 tumor types), led by canonical oncogenes *EGFR* and *SRC*. Negative regulation of anoikis (23 cancers elevated), led by amplifications in caveolin (*CAV1*), *SRC*, *PIK3CA*, and FAK/*PTK2* (**Figure 3B**) were the third highest. Among the most commonly altered was keratinization, which drives cell cycle progression in breast epithelial cells (Sack et al., 2018). Other frequently elevated pathways include amoebiasis, cAMP signaling, DNA methylation, and female meiotic division.

**Figure 3.**
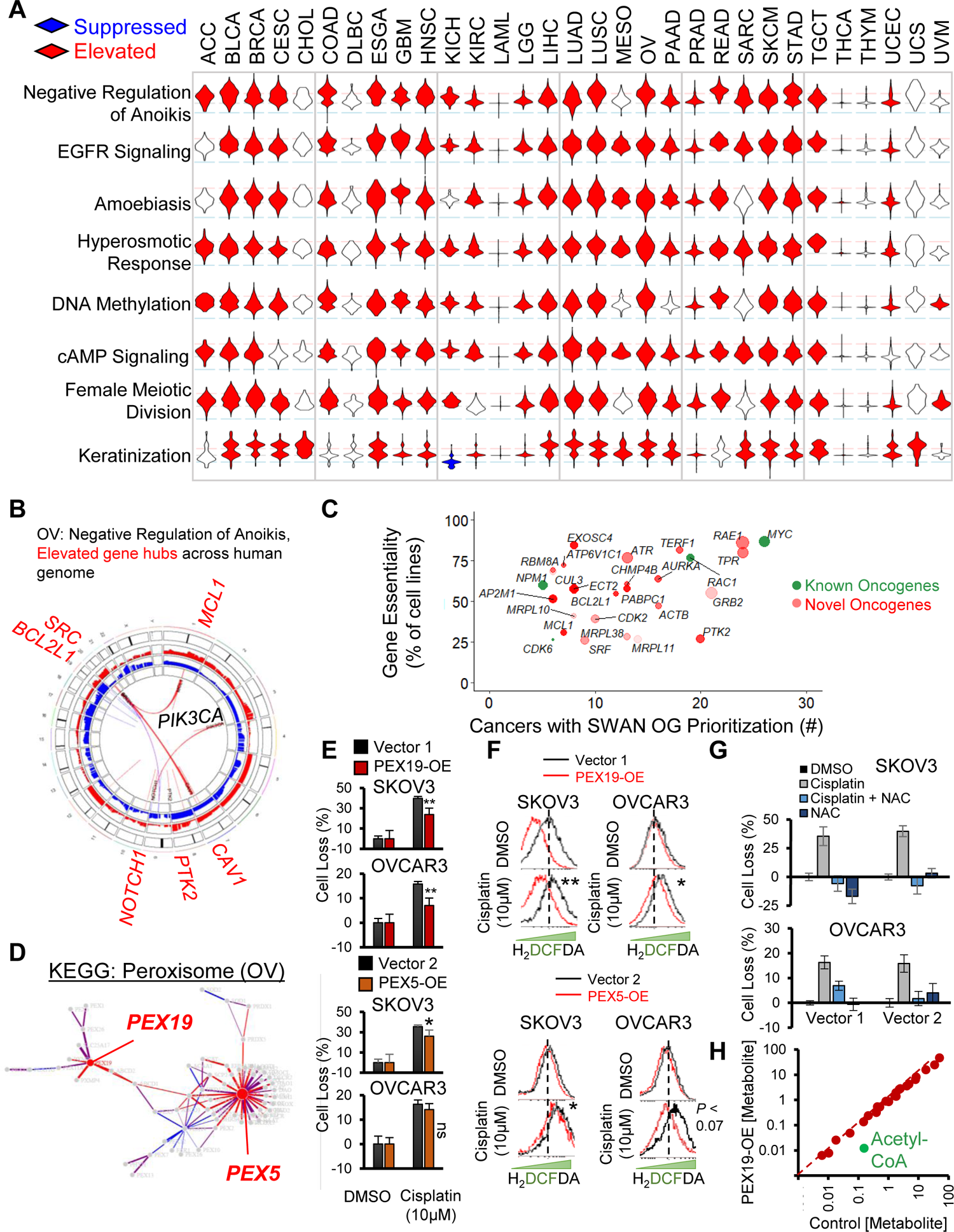
Pan-cancer elevated CNA pathways. (**A**) Unusually pervasive elevated CNA pathways. Violin histograms of SWAN scores with red fill indicating significant (FDR < 0.0001) pathway elevation and blue fill indicating significant pathway suppression. (**B**) SWAN Circos plot. Red and blue outer rings are frequency plots of gains or deletions and the inner ribbons represent genes within the selected pathway. Labeled gene symbols are enriched for CNA gains. (**C**) Impact summary of novel pan-cancer SWAN elevation-prioritized genes. Green color indicates COSMIC OGs. Size is proportional to the mean z-score SWAN contribution across cancers with OG-containing pathway elevation. Higher transparency indicates interacting protein genes influenced each gene’s identification by SWAN, rather than CNAs of the gene itself. (**D**) SWAN network generated, with edges represent protein-protein interactions. Blue nodes are enriched for loss CNAs and red nodes (such as PEX5 and PEX19) are enriched for gain CNAs. (**E**) Mean ± standard error of crystal violet viability assays comparing 48h cisplatin to control 0.1% DMSO treatment. N = 4 experiments with data combined from all experiments. (**F**) Flow cytometry of ROS indicator H_2_DCFDA following 48h cisplatin or control 0.1% DMSO treatment in PEX19 overexpressing or PEX5 overexpressing cells. Significance determined from N = 3 independent experiments. Dotted line marks peak control stain. (**G**) Mean ± standard error of crystal violet viability assays comparing 48h cisplatin to control 0.1% DMSO treatment with or without 2mM NAC. N = 2 experiments. (**H**) Summary of metabolite concentrations within a metabolomic study comparing N = 5 PEX19 overexpressing SKOV3 cells relative to control vector cells. Acetyl-CoA is highlighted as an outlier. \**P* < 0.05, \*\**P* < 0.01, ns *P* > 0.05.

Modern cancer therapies are often developed to target genes that are overexpressed or constitutively active in cancer. To evaluate novel CNA targets, we compared SWAN interactome prioritization data with sgRNA screens of 324 cancer cell lines (Behan et al., 2019). **Figure 3C** depicts putative (defined here as not Tier 1 COSMIC) OGs which were identified as a dependency gene in at least 25% of cancer cell lines (**Table S3**). Sixty-five additional prioritized OG nodes were found which were not analyzed in the 324-cell line screen (**Figure S6A, Table S3**). Included within these putative novel OG CNAs are emergent targets for cancer therapy. Of note, *ADORA2A* encodes adenosine receptor A2a, which negatively regulates inflammatory immune response (Ohta et al., 2006), and its blockade enhances pre-clinical syngeneic models of PD-1, TIM-3, or CTLA-4 therapies (Leone and Emens, 2018). Another, *PTK2*, encodes focal adhesion kinase (FAK), which enables cells to survive a loss of adhesion (Sulzmaier et al., 2014). Two Phase II oncology trials target FAK with a small molecule inhibitor defactinib (NCT02465060 and NCT03727880). Future studies may consider the SWAN prioritized CNA-altered OGs as therapeutic targets.

Some pathway disruptions were unique to a single or a handful of cancers. Such pathways may represent unusually selective pathways for targeted treatment or early diagnosis. One largely unexplored but selectively elevated pathway involved peroxisome biogenesis. Second to testicular cancer, OV was most affected by pathway CNA elevation (**Figure S6B**), but not by mutation (**Figure S6C**). SWAN networks highlighted *PEX5* and *PEX19* as the most relevant amplified genes in serous ovarian cancer (**Figure 3D, Figure S6D**). *PEX5* is within an elevated CNA in 53.6% of OV tumors and *PEX19* in 57.8% of OV tumors (**Figure S6E-G**). *PEX5* and *PEX19* function to properly import peroxisome membrane proteins to the organelle (Kim and Hettema, 2015). Overexpression (-OE) of either *PEX5* or *PEX19* modestly reduced cisplatin lethality within SKOV3 and OVCAR3 ovarian cancer cells (**Figure 3E**), neither of which contain amplifications of either gene (**Figure S6H**). To investigate if this phenotype was due to a reduction of intracellular reactive oxygen species (ROS), which peroxisome metabolism contributes to (Schrader and Fahimi, 2006), we stained cells for cisplatin-stimulated ROS. PEX5-OE or PEX19-OE reduced the amount of cisplatin-stimulated ROS (**Figure 3F**). Scavenging ROS with N-acetyl cysteine (NAC) similarly abrogated cisplatin toxicity in the control cell lines (**Figure 3G**). These results were not initially expected since peroxisomes can be a source of ROS, primarily through catabolic Acyl-CoA oxidase function for lipid β-oxidation (Poirier et al., 2006). To test directly if lipid metabolism was disrupted, we performed unbiased ultra-performance liquid chromatography mass-spectrometry metabolomics. An outlier metabolite in PEX19-OE SKOV3 cells was a 92% reduction in Acetyl-CoA, a product of β-oxidation (**Figure 3H**, *P* < 0.051). Overall lipid content of the cells was unchanged (**Figure S6I** and **Table S4**), suggesting that PEX19-OE cells may need to replace Acetyl-CoA via exogenous sources of lipids, a known phenotype of ovarian cancer (Motohara et al., 2019). *PEX5* expression correlated with poor prognosis, whereas *PEX19* did not (**Figure S6J** and **S6K**). In summary, SWAN identified two potential OGs within the peroxisome KEGG pathway, and overexpression of each was sufficient to reduce ROS generation in chemotherapy-stressed ovarian cancer cells.

### Identification of known and novel tumor suppressor pathways

Focal deletion regions of *PPP2R2A*, *CDKN2A*, *ATM*, *NOTCH1*, *TP53*, *PTEN*, and the *BRCA1/2* genes have been previously highlighted (Zack et al., 2013). In our pan-cancer analysis, these tumor suppressors were often highly influential in determining suppression of pathways; *PTEN*, *TP53*, and *BRCA1/2* are highlighted in the top 1% of suppressed pathways (**Table S2**). Phospholipid dephosphorylation was the most commonly suppressed pathway, found as significantly suppressed in 22 of the 31 tumor types studied (**Figure 4A**). Along with *PTEN*, losses in phosphatidylinositol 4,5-bisphosphate 5-phosphatases (*INPP5E/J/K*), lipid phosphate phosphohydrolase (*PPAP2A/C*), lipid phosphate phosphatase-related protein (*LPPR1/3/4*), and synaptic Inositol 1,4,5-Trisphosphate 5-Phosphatase (*SYNJ2*) were commonly dysregulated by deletions. Replicative senescence (**Figure 4B**) and apoptotic regulators (**Figure 4C**) are suppressed as expected. Phospho-STAT signaling was suppressed in 17 tumor types, led by deletions in Type I interferon genes. Attachment of spindle microtubules to the kinetochore was suppressed in 20 tumor types, led by deletions connected to Aurora kinases (*AURKB/C*), kinetochore microtubule motor *CENPE*, and anaphase promoting complex regulator *BUB3*. OV, a highly aneuploid tumor type, was amongst the most suppressed for protein localization to the kinetochore, due to distributed losses on Chr1p, Chr15q, and Chr17. Chromatin organization was suppressed in 21 cancer types, with deletions in p53, *ATM*, β-catenin, sterol regulatory element-binding transcription factor 1 (*SREBF1*), lysine (K)-specific demethylase 1A (*KDM1A*), and histone acetyltransferase p300 (**Table S2**). The most commonly identified TSGs were amongst the best-established tumor suppressors. SWAN interactome analysis predicts p53 as the most significant and common TSG (**Figure 4D**). Altogether, 170 novel TSGs were identified (**Table S3**).

**Figure 4.**
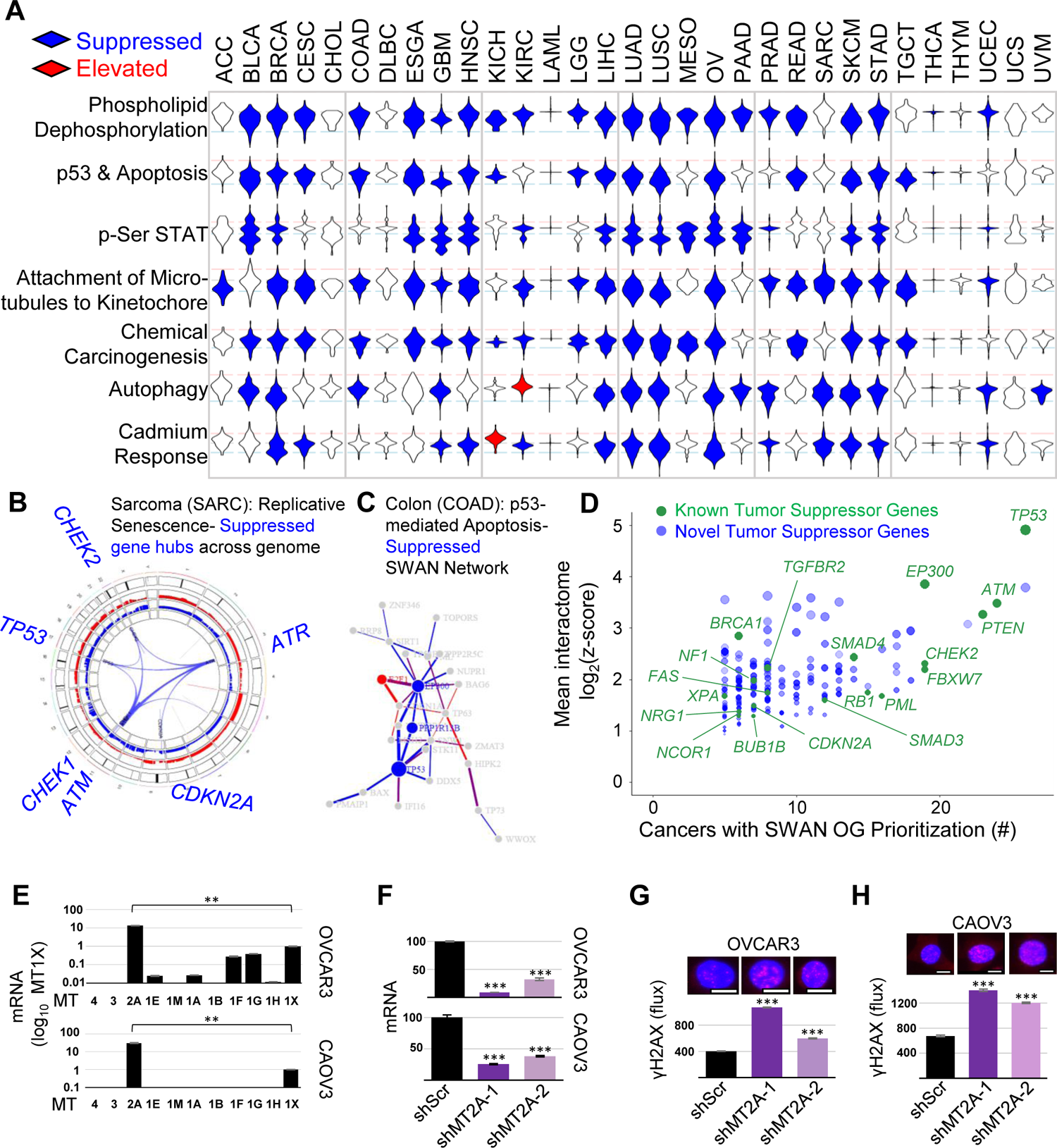
Pan-cancer suppressed CNA pathways. (**A**) Unusually pervasive suppressed CNA pathways. Violin histograms of SWAN scores with blue fill indicating significant (FDR < 0.0001) pathway suppression and red fill indicating significant pathway elevation. (**B**) SWAN Circos plot. Red and blue outer rings are frequency plots of gains or deletions and the inner ribbons represent genes within the selected pathway. Labeled gene symbols are enriched for CNA losses. (**C**) SWAN network generated, with edges represent protein-protein interactions. Blue nodes are enriched for loss CNAs and red nodes are enriched for gain CNAs. (**D**) Known and novel TSGs discovered by interactome analysis of all 31 cancer types analyzed, with those present in at least 5 tumor types displayed against z-score values. Green color indicates previously known COSMIC TSGs. Size is proportional to the mean z-score SWAN contribution across cancers with TSG-containing pathway suppression. Higher transparency indicates interacting protein genes influenced each gene’s identification by SWAN, rather than CNAs of the gene itself. (**E**) RT-qPCR data of metallothioneins within 16q gene cluster. (**F**) Validation of shRNA-mediated knockdown of MT2A by RT-qPCR. (**G**) Genotoxic damage as measured by γH2AX immunofluorescence in the presence of 100 µM cadmium is shown for OVCAR3 cells and (**H**) in the presence of 50 µM cadmium for CAOV3 cells. (I-L) N = 3 experiments, with mean ± s.e.m. shown.

Protein quality control and cellular homeostasis was commonly disrupted across cancers. Protein deglycosylation was suppressed (19 of 31 tumor types) most commonly through deletions related to *EDEM1* and *DERL2*, proteins involved in extracting misfolded glycoproteins as part of endoplasmic reticulum associated degradation. Protein processing in the endoplasmic reticulum was a commonly suppressed KEGG pathway (14 of 31 tumor types). We previously found autophagy, by *MAP1LC3B* and *BECN1* gene loss, to be suppressed and therapeutically targetable in OV (Delaney et al., 2017). *BECN1* is a bona-fide tumor suppressor in ovarian cancer and contributes to genome instability (Delaney et al., 2020; Kumar et al., 2020a). In this pan-cancer analysis, autophagy was suppressed in many other tumor types as well (14 of 31 tumor types), as was protein ubiquitination and degradation (9-14 tumor types). Ion homeostasis was commonly disrupted; negative regulation of potassium transport (9 tumor types), manganese ion transport (8 tumor types), copper ion homeostasis (7 tumor types), and zinc or cadmium ion response (10 or 15 tumor types, respectively) were pervasively suppressed by CNAs.

To confirm if SWAN identified novel tumor suppressor pathways which are biologically relevant, we selected the zinc and cadmium response pathways. These are dominated by concomitant loss of metallothionein genes in a cluster on Chr16q. Ovarian tumors lose this gene cluster in 60% of high-grade serous tumors. Metallothioneins are cysteine-rich proteins which chelate divalent cations within the cell: particularly Zn^2+^ and toxic heavy metals such as Cd^2+^ (Klaassen et al., 1999). Cadmium is an environmental toxin thought to increase lung, pancreatic, and endometrial cancer risk (McElroy et al., 2017). Using ovarian cancer cell lines, metallothionein-2A (*MT2A)* mRNA was most highly expressed amongst all isoforms (**Figure 4E**). Therefore, *MT2A* was knocked down in ovarian cancer cells (**Figure 4F**). Loss of the metallothionein gene cluster may contribute to cadmium-mediated oncogenesis by allowing for genomic instability. To test this hypothesis, knockdown cells were evaluated for γH2AX foci following cadmium exposure. *MT2A* knockdown cells contained more γH2AX foci than control cells (**Figure 4G** and **4H**), consistent with the hypothesis that these metallothionein genes protect cells against cadmium-mediated genotoxic damage.

### Cancer-specific pathway alterations

Dysregulated pathways in distinct cancer types may be particularly informative. Glioblastoma multiforme (GBM) is the only tumor type elevated in “positive regulation of neuron death,” while 19 tumor types are haploinsufficient (**Figure 5A**). This was due to *SRPK2* elevation, which has recently been implicated in RNA dysregulation in GBM (Song et al., 2019) via phosphorylation of SRSF3 (Long et al., 2019), and elevation of *PTPRZ1*, which macrophages stimulate for GBM stem cell growth (Shi et al., 2017), *CDK5*, which is involved in neuronal migration, and canonical oncogenes *MAP2K7* and *ABL1* (**Figure 5B**). GBM cells may alter this pathway in a single chromosome gain event, as *SRPK2*, *CDK5*, and *PTPRZ1* are all encoded on Chr7q (**Figure 5C**). By CAIRN analysis of CNAs (Delaney et al., 2020), these genes are co-incidentally gained in 45% of GBM tumors.

**Figure 5.**
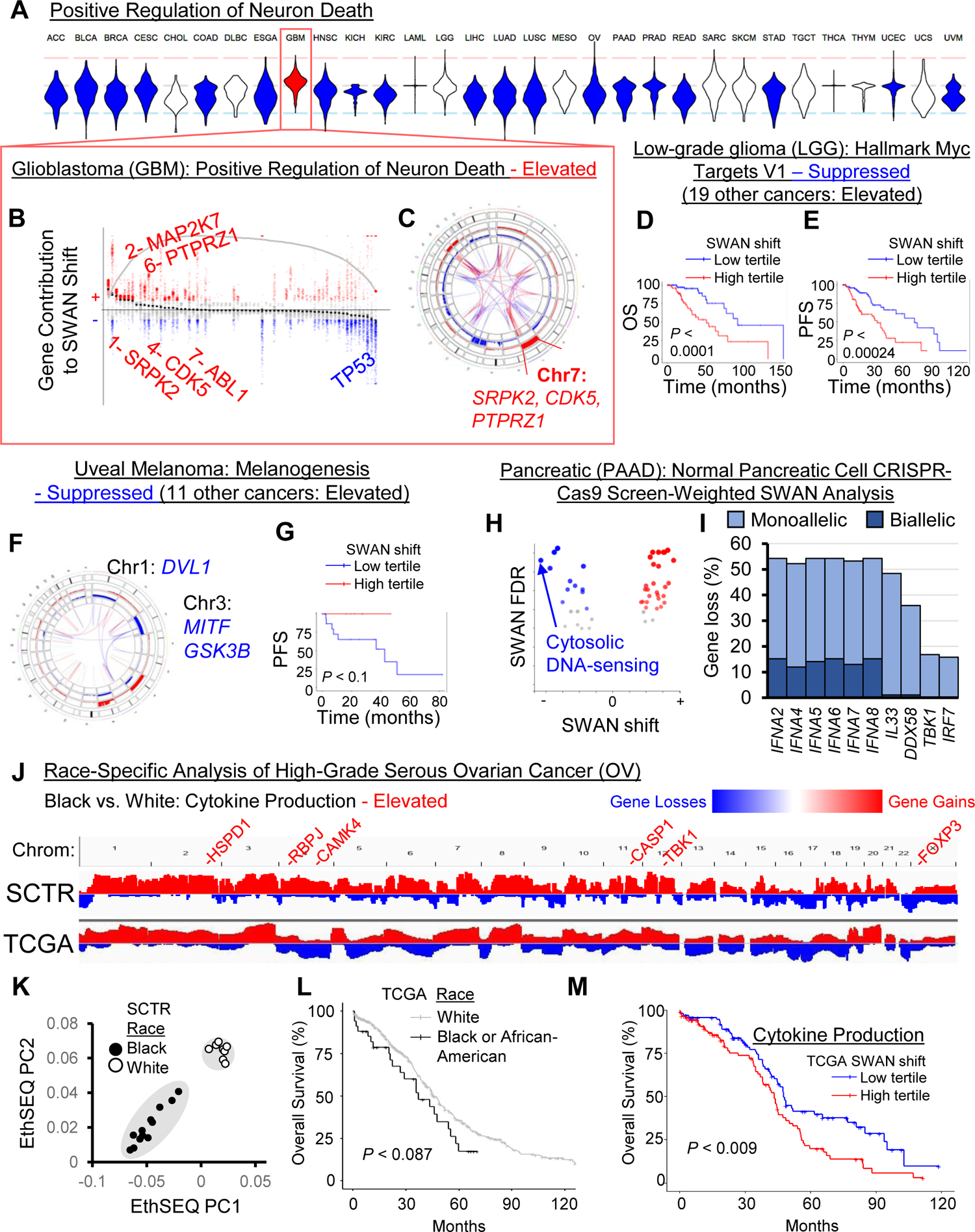
Cancer-specific CNA patterns. (**A**) Rare pathways had opposite SWAN shifts relative to the majority of cancer types. Shown is the example of “GO: Positive regulation of neuron death” which was uniquely upregulated in the brain cancer GBM, as illustrated in a (**B**) SWAN feather plot and (**C**) Circos plot. (**D**) Overall survival (OS), *P* is from Kaplan-Meier analysis. (**E**) Progression-free survival (PFS) plot. (**F**) SWAN Circos plot. (**G**) Progression-free survival (PFS) plot. (**H**) SWAN analysis scored by HPNE CRISPR-proliferation screen hits in place of haploinsufficiency. The most suppressed pathway by magnitude is highlighted. (**I**) The most frequently deleted genes from (h) pathway. (**J**) Integrative Genomics Viewer cohort summary plots for the new African-American enriched OV cohort (SCTR) compared to The Cancer Genome Atlas (TCGA) OV cohort. Noted genes indicate SWAN prioritized genes within indicated pathway. (**K**) EthSEQ analysis and principal component clustering of variants in the SCTR cohort. Self-identified race is labeled for black and white patients. (**L**) Kaplan-Meier analysis of TCGA tumors separated by self-identified race and plotted for overall survival. (**M**) Kaplan-Meier plot of TCGA data separated by SWAN shifts.

In Low-Grade Glioma (LGG), Hallmark Myc Targets V1 was marked as “suppressed” whereas 19 cancer types were marked as “elevated”. Interestingly, the subset of tumors with CNA losses within MYC targets both had greater overall survival (**Figure 5D**) and progression free survival (**Figure 5E**) in LGG. These genes are enriched on Chr1p and Chr19q, which was identified in the TCGA publication as prognostic of *IDH1* mutant tumors (Cancer Genome Atlas Research et al., 2015). LGG is often driven by *MYC* or *IDH1/2* mutations, suggesting that the tumors which have spontaneously lost an array of MYC targets may have attenuated their oncogenic potential.

Uveal melanoma (UVM) exhibits an unusual mode of initial extravasation which first involves intercalation with endothelial cells (Onken et al., 2014). While 11 cancers are elevated in the KEGG pathway melanogenesis, which usually involves gains in Wnt-β-catenin regulating factors *FZD1*, *GNAI1*, and *WNT3A*, UVM was the sole cancer suppressed in this pathway due to Chr1 and Chr3 losses overlapping *MITF*, *GSK3B*, and *DVL1* (**Figure 5F**). Patients with these losses are in the poor prognosis group, particularly for progression free survival (**Figure 5G**). Poor prognosis is associated with increased immunosuppressive profiles (Figueiredo et al., 2020).

To address tissue specificity in an unbiased fashion, the top 1% variable pathways were K-means clustered. There were clearly different cancer subsets with regard to phospho-STAT signaling, keratinization, epigenetic regulation of rDNA, protein carboxylation, and serine peptidase (**Figure S7A**). Cancer types did not strongly cluster together. The remaining cancer-specific altered pathways can be found in **Table S2**.

### Tissue-specific annotation weighting reveals a suppression of cytosolic DNA response

Whole-genome CRISPR-Cas9 screens of non-transformed normal tissue have identified tissue-specific drivers of proliferation (Sack et al., 2018). In lieu of haploinsufficiency data, we applied weights on pancreatic cancer (PAAD) CNAs in SWAN using genes enriched for proliferation changes from a CRISPR-Cas9 screen in primary immortalized pancreatic HPNE cells. In a KEGG pathway analysis, the cytosolic DNA-sensing pathway was the most suppressed pathway in PAAD (−9.4 SWAN shift, FDR ≤ 8.1×10^-8^, **Figure 5H**). This was led by the *IFNA* genes, which produce type I interferons and act as positive feedback inducers of a central dsDNA sensor cGAS (Ma et al., 2015) (**Figure 5I**). Another suppressed gene was *DDX58*, a primarily dsRNA sensor which can also detect some types of dsDNA (Zevini et al., 2017). Chromosome instability often results in micronuclei, which normally activate cGAS-STING signaling. However, in some cancer cells this pathway was found to be attenuated by an unknown mechanism (Bakhoum et al., 2018) allowing for cell survival and increased metastasis. CNAs may be one mechanism cancer cells use to reduce cGAS-STING pathway signaling, particularly in pancreatic cancer. Weighting by tissue-specific sgRNA screens thus yielded further insights into CNA patterns and tumor biology.

### SWAN case study on race-specific CNA patterns

African American data represents only 6% of the tumors present in TCGA OV data. Expansion of data and analysis in this group is warranted. We obtained 12 tumors from African American high-grade serous ovarian cancer patients and 8 non-Hispanic white patient controls and performed whole-exome sequencing (**Table S5**). Since normal tissue was not available for these unique samples, confident somatic SNV analysis was complicated by rare but normal variants. CNAs, conversely, are uncommon in normal tissue and ascertainment of CNAs was possible using Control-FREEC software (Boeva et al., 2012). This method was remarkably similar in overall cohort CNA calls to TCGA analyzed tumors (**Figure 5J**). To investigate the possible race-specific CNAs in African American patients, SWAN was used to compare African American patients to white patients from this study as well as combined with the TCGA study. The cytokine production pathway was found to be significantly elevated in African American patient tumors relative to white patient tumors (**Figure 5J** gene labels). Self-reported race matched race-defining variants found by EthSEQ analysis on these tumors, with expected higher admixture present in African American patients (**Figure 5K**). Black OV patients respond poorly to therapy relative to white patients, even when taking into account socioeconomic factors and comorbidities (Hildebrand et al., 2019). This trend, albeit not significant, was seen in TCGA survival data (**Figure 5L**). SWAN shifts mapping to elevation of cytokine production were associated with poor prognosis in OV overall (**Figure 5M**). Existing socioeconomic factors which lead to persistent inflammation in black patients may allow for de-repressed cytokine production in black patient tumor cells, which would otherwise allow T-cell responses to clear tumors. Low-dose rapamycin treatment may re-enable T-cell clearance within these patients (Mannick et al., 2014; Mannick et al., 2018).

### Pathways associated with SNV mutant drivers

Each SNV/indel driver mutation may be predicted to require its own set of CNAs to assist in cancer development. To test this hypothesis, we analyzed which tumor types had CNA-altered pathways within the subset of specific mutant tumors, relative to non-mutant tumors of the same histotype. We tested commonly mutated TSGs/OGs: *TP53*, *CDKN2A*, *KRAS* or *NRAS* or *HRAS*, *BRAF*, *BRCA1* or *BRCA2*, *EGFR*, *PTEN*, *HIF1A* or *VHL*, *RB1*, *ATM*, *APC*, and *MSH*s (*MSH2,3,4,5* or *6*). Of all of these possible driver mutations, *TP53* had the most pathways commonly affected in multiple cancers (**Table S6**). This is consistent with the observation that *TP53* mutation is the most significantly associated with aneuploidy by multiple orders of magnitude (Taylor et al., 2018). A suppression of KEGG: autophagy in p53 mutant subsets of tumors was observed in 8 cancer types (**Figure 6A**). Uterine/endometrial cancer is known to have worse prognosis with p53 mutation, and these tumors were severely reduced in autophagy gene content (**Figure 6B**). OV, the cancer type we have thoroughly investigated for its loss of autophagy (Delaney et al., 2020; Delaney et al., 2017; Kumar et al., 2020a), was not found in this set due to the ubiquity of p53 mutations, precluding a non-mutant control comparison. Mutation in p53 is commonly associated with elevation of GO: Regulation of Cell Adhesion Mediated by Integrin (14 tumor types elevated, led by FAK/*PTK2* and *LYN*).

**Figure 6.**
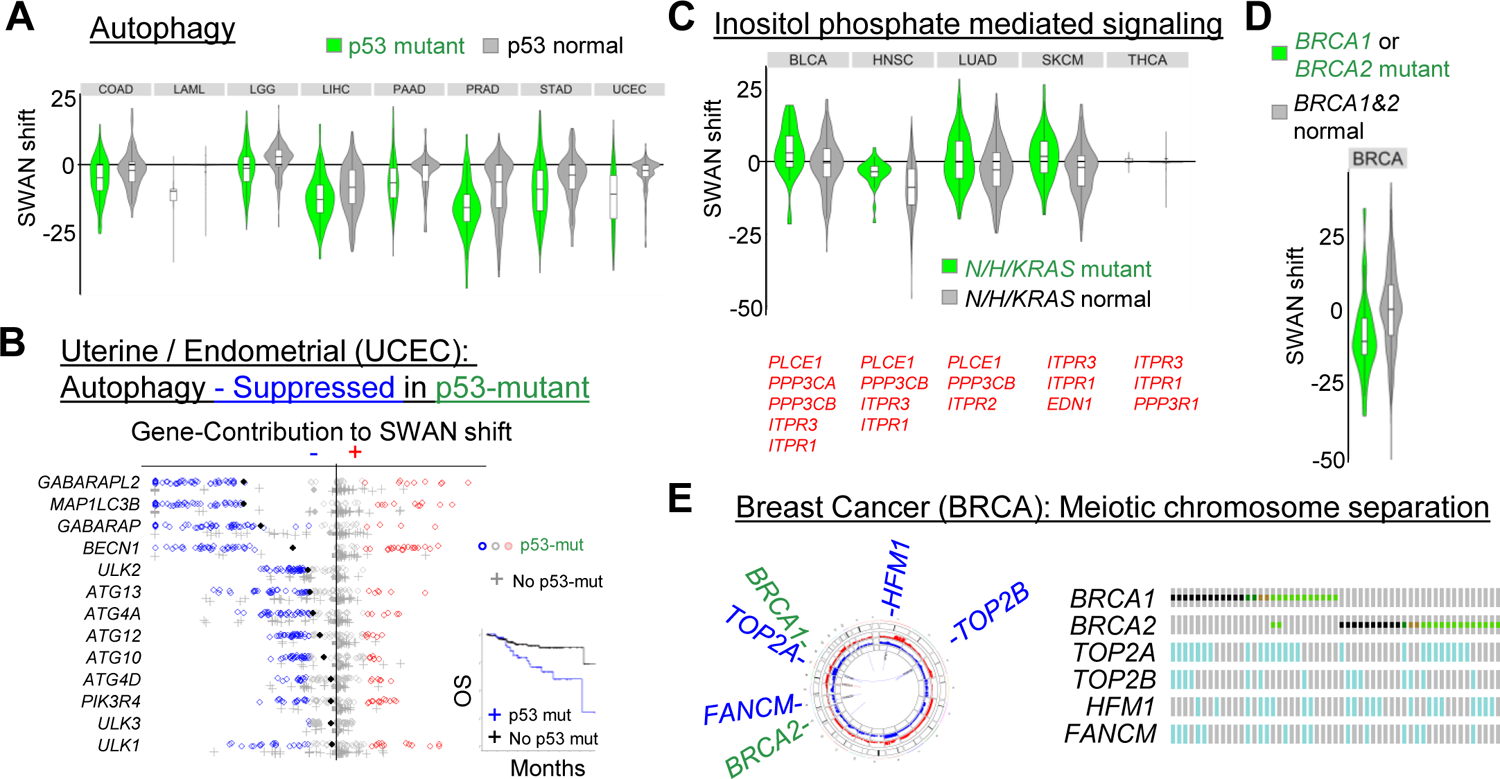
Mutation associated SWAN shifts. (**A**) All cancers with a significantly different SWAN shift spectrum for p53 mutant tumors compared to p53 wild-type tumors are shown for the KEGG: Autophagy pathway. (**B**) SWAN feather plot of suppressed autophagy genes. UCEC samples with p53 mutation are plotted as circles, samples with no p53 mutation as crosses. Filled diamonds represent mean SWAN shifts per gene. Inset panel shows overall survival of p53 mutant UCEC compared to non-mutant UCEC. (**C**) All cancers with a significantly different SWAN shift spectrum for RAS (*NRAS*, *HRAS*, or *KRAS*) mutant tumors compared to RAS wild-type tumors are shown for the GO: Inositol phosphate mediated signaling pathway. (**D**) Breast was the only cancer with a significantly different SWAN shift spectrum for *BRCA1* or *BRCA2* mutant tumors compared to BRCA wild-type tumors for the GO: meiotic chromosome separation pathway, with (**E**) pathway genes highlighted for regions of gene deletion by Circos plot (left) or Oncoprint (right).

The next most consistently altered pathway sets occurred in RAS mutant cancers. In these cases, the commonly dysregulated pathways may further enhance the activation of the RAS pathway. RAS mutation was associated with an increase in inositol phosphate mediated signaling (**Figure 6C**), likely increasing activation of PI3K (Castellano and Downward, 2011). The remainder of known, common driver mutations were distinct to individual cancer types. For example, the *BRCA1/2* driven cancers were associated with a decrease in meiotic chromosome condensation specifically in breast cancer (**Figure 6D**). This pathway was suppressed via allelic loss in *FANCM*, *TOP2A/B*, and *HFM1* (**Figure 6E**). Since CNAs are far more common than individual driver SNV mutations, more samples are needed to provide the statistical threshold for CNA pathway differences associated with other individually rare SNV drivers. Overall, these data support the model that SNV/indel mutations alter biology in a manner which is further exacerbated by CNAs.

### Prognostic alterations

If SWAN-identified CNA pathways drive the biology of tumors, then it would be expected that some pathways influence patient prognosis. Both cBioPortal and ProteinAtlas offer survival curves comparing “low” and “high” expression of individual genes for this purpose. Here, we explored whether pathway scores can separate prognostic groups. In a log-rank test analysis using whole-cohort data, 12,781 pathways from 18 cancers were found to be significantly prognostic by comparing upper and lower tertiles of SWAN shifts. Survival estimates can be misleading (Crespo et al., 2016), and we accordingly found an ability to erroneously call a single pathway as significantly positively prognostic or negatively prognostic, depending on which patients were randomly selected (**Figure 7A, Figure S7B**). Desiring to prioritize the 12,781 pathway hits into those most likely to be medically informative, we developed a machine learning approach. A random subset of two-thirds of tumors was used to build Cox-proportional hazard (Cox-PH) models from SWAN network shift data. The remaining one-third of tumors per random selection was used to generate hazard ratios using the Cox-PH model. 101 random patient selections were used to determine which pathways scored as similarly prognostic in the test set. While Cox-PH is traditionally used for multiple covariates, its predictive capacity for single variables was herein used. The machine-learning approach here required an identical hazard ratio direction and subsequent test group log-rank significance in >80% of patient selections to call a pathway as prognostic (**Figure 7B**). This strategy was amenable to 18 cancer types with sufficient survival data and reduced the prognostic pathways from 12,781 pathways within the pan-cancer cohort to 1,696 pathways (**Figure S7C-S7E)**. 1,079 are from the highly predictable LGG data, leaving 617 machine-learned prognostic pathways from 17 cancer types. While an ideal machine-learning approach would include an independent dataset, data of sufficient size and comparable form across cancer types were unavailable. Our approach nonetheless sharply narrowed the scope of likely prognostic CNA pathways. Kaplan-Meier curves of the entire cohort confirm that machine learned pathways were negatively prognostic (**Figure 7B**) and positively prognostic (**Figure 7C**), suggesting that SWAN data may be used to categorize patients by prognosis. Machine-learned prognostic pathways for each cancer type are supplied as **Table S7**.

**Figure 7.**
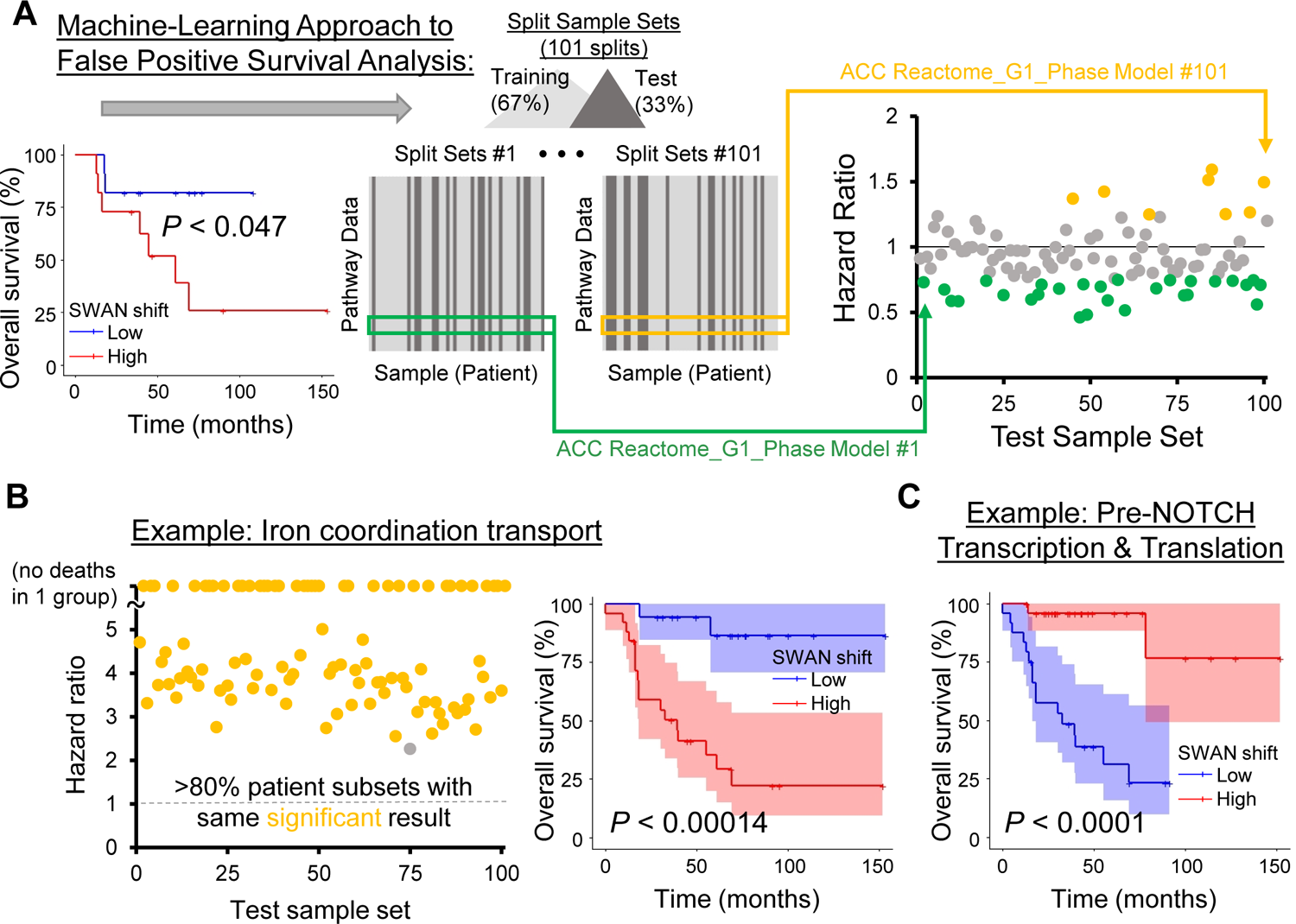
Machine-learning approach for improved prognostic estimates of CNA influenced pathways. (**A**) An example of a nominally-significant prognostic pathway, corrected by a machine learning approach. Patient data are first separated by SWAN pathway shifts, with the “low” and “high” groups separated by 1 SD centered at the median shift. With the entire patient cohort considered, a Kaplan-Meier analysis yields *P* < 0.047. 101 training sets of SWAN shift data build risk scores, then the upper quartile risk is compared to the lower quartile risk group by Kaplan-Meier analysis to yield a *P*-value for the test group. Colored circles indicate HR at least 1 SD from the null 1 value. Potential false positive prognostic pathways were removed by (1) determining significance in the applied CoxPH model risk for each test set, and (2) determining CoxPH HR direction in each training set. The most prognostic pathways are labeled as those with >80% of randomizations as both significant and in same HR direction, such as in example (**B**). A Kaplan-Meier overall survival curve with 95% confidence interval is shown for the whole patient cohort (right panel). (**C**) A negatively prognostic Kaplan-Meier overall survival curve with 95% confidence interval is shown. In this case, 98% of patient picks were significant.

### Online atlas

We uploaded a SWAN analysis of 31 cancer types studied by TCGA into an online portal. The accessible data include SWAN CNA network-level scores and per-patient scores for all cancer types for four pathway sets: GO, KEGG, Hallmark, and Reactome. This database is provided online at https://delaney.shinyapps.io/cnalysis_tcga_atlas/. Summary data for all analyses discussed here are also provided as **Tables S1-S7**.

## Discussion

Copy-number alterations represent a wealth of unexplored oncogene and tumor suppressor gene data within the cancer genome landscape. Expanse and heterogeneity in these data have previously prevented identification of biologically relevant CNAs. SWAN demonstrably improves the identification of TSGs and OGs on CNAs within a pan-cancer dataset. In addition to this computational validation, wet-lab validation was performed. SWAN identified *PEX5* and *PEX19* as elevated genes within the peroxisome biogenesis pathway. These genes each conferred platinum resistance by oxidative control. SWAN marked cadmium response as suppressed in 15 tumor types, and knockdown of metallothionein 2A conferred increased γH2AX foci formation in response to cadmium. Newly sequenced tumors analyzed by SWAN determined that elevation of cytokine production genes occurs within tumors from an underrepresented racial group. SWAN was developed with an emphasis on ease-of-use and widespread applicability so that SWAN may be utilized in future experiments with complex genetics. While optimized for oncology, SWAN may be considered for use as generally applicable pathway analysis software.

A limitation with modern genetically-targeted cancer treatments is the small percentage of patients who may benefit from a specific genetic alteration. Some estimates are that only 8.3% of patients are eligible for genetically-targeted therapy (2018), compared to 5.1% over a decade prior (2006) (Prasad, 2020). In OV, which lacks canonically targetable mutant drivers, alternative tests have been developed to test for functional deficiency in homology directed repair (HDR), thereby enabling PARP inhibitor targeted therapy even in the absence of *BRCA1/2* mutations (Longo, 2019). This HDR deficiency is influenced by the ubiquitous CNAs described here. Our analysis of CNAs represents a wealth of potentially targetable pathways that are altered in double-digit percentages of patients. Moreover, therapeutic windows as a result of tumor CNAs are of high likelihood since functional normal cells do not contain clonal somatic CNAs. Targeting pathways, rather than individual genes, may reduce the likelihood of resistance mutation development and resurgent cancer, especially if multiple drugs are used (Bozic et al., 2013).

While previous CNA studies have thoroughly studied focal homozygous deletions and arm-level aneuploidy in cancer, these studies do not provide a comprehensive statistical assessment of what patterns of changes can alter biological pathways. This is critical because a pathway may be suppressed via multiple chromosome arms or monoallelic changes which may differ tumor-to-tumor. An example of this is the keratinization pathway. While it was previously shown that single gene overexpression of keratinization factors can promote cell cycle progression, these effects were cell type specific (Sack et al., 2018). However, those cell-type specificity results must be a consequence of other factors within the cell which ameliorate or exacerbate the effects of single gene overexpression. Networks, conversely, take into account these disparate interacting factors that differ between individual tumor samples. An analysis of focal-amplification regions consistently altered across tumors identified regulators of Nf-κB, Wnt/β-catenin, *MYC*, *AKT*, *ERBB2*, Cyclin-D1 and -E1, and *TERC* (Zack et al., 2013). Pathway alterations within these commonly amplified segments centered on chromatin modifiers such as *BRD4*, *KDM2A*, and *KDM5A* (Zack et al., 2013). These genes are well-known oncogenes and were often identified as impactful by SWAN. However, SWAN was able to identify cancer addictions depleted in sgRNA screens and further prioritized 65 novel oncogenes. Pan-cancer and cancer-specific CNA pathways were identified and their relevance to primary chemotherapy can be referenced to the machine-learned prognostic pathways released here.

## STAR Methods

### Key Resources Table

See supplement

## Experimental Model and Subject Details

### Cell Lines

Established cell lines were purchased from the American Type Culture Collection (ATCC) and validated by short tandem repeat (STR) profiling (Duke University and ATCC).

### Human High-Grade Serous Ovarian Cancer Samples

Flash frozen samples were requested from biorepositories. All samples were stage 2C or higher high-grade serous fallopian or ovarian cancer, with the exception of a single control Caucasian sample with paired uterus normal control used for bioinformatic quality control. Cancer stage, self-reported race and ethnicity (either non-Hispanic black or non-Hispanic white), and age at diagnosis is reported in **Table S5**.

## Method Details

### Core algorithm of SWAN

#### SWAN Inputs

SWAN takes as a data input any matrix of samples in columns and annotations in rows. For the examples here, tumors from individual patients were columns, and genes were set in rows. However, in principle any numeric set of data may be used. Aside from this primary experimental data, a secondary identical-format control set of data may be input. Sample sizes may differ between experimental and control data. To create and score networks, additional optional inputs include: haploinsufficiency data (yeast and mouse homolog data (Delaney et al., 2017), by default), interaction data (protein-protein interaction data from BioGRID (Stark et al., 2006) by default), mutation data, and RNA data. Pathway sets need to be defined, and users may create and upload pathways for user-specific annotations. Default pathways sets are from MSigDb (Liberzon et al., 2011) and are gene-centric. Tuning inputs include: number of permutations to perform (default: 200), a vote-factor (default: 25, which assists in *P*-value stability), a maximum and minimum number of genes within a pathway to be tested (default: 10-200 genes, to avoid large hairballs), and toggles for choosing to score or not score any of the above.

For the analyses presented here, the matrix input was gene-level copy number alterations. The algorithm is optimized for copy-number alteration data with integer values (−2 for 0N, −1 for 1N, 0 for 2N, 1 for 3N, 2 for 4N or more). TCGA data were downloaded from cBioPortal using “Provisional” datasets (Gao et al., 2013). Mutation data included TCGA mutation data with a “1” marking a non-synonymous mutation and a “0” for no non-synonymous somatic mutation. RNA data were log2 per-gene normalized TCGA microarray data from the pan-cancer TCGA set, “EBPlusPlusAdjustPANCAN_IlluminaHiSeq_ RNASeqV2.geneExp.tsv” (Group et al., 2020). Pathway sets used were the Hallmark, KEGG, Reactome, and GO.

#### Data structure considerations for input SWAN data

Raw input data should be normalized in a reasonable fashion to ensure the range of values is roughly symmetric around the median. That is, if extremely positive or negative values are rare but present, they should be capped at a consistent value (eg, for copy-number analysis, 200N is capped at 4N). If data are not normalized or thresholded, then the deviations between permutations become enormous and statistical significance suffers. If choosing to include RNA with DNA analysis, RNA data should be normalized to have the same range (−2 to +2). Convenience normalization functions are provided in the R package, and can be used in the online tool by marking a check-box. If choosing to include mutation data, non-mutated genes should be “0”, mutant genes should be “1”. If a user chooses to prioritize some genes’ mutations over others, a custom scoring multiplier per gene may be uploaded to replace “haploinsufficiency” scoring information, or the “1” value may be modified to any other positive integer (skewing from the median for mutations is acceptable).

Some users may prefer directional networks. To perform this type of analysis, users need to create a separate pathway set containing a pair of pathways for each direction: (a) genes that positively alter the pathway and (b) genes which negatively alter the pathway. An example is the MSigDB GO Biological Process pathway set, limited to “GO_POSITIVE_REGULATION_” and “GO_NEGATIVE_REGULATION_” pathways. Since this is conceptually easier to interpret the output data, we have included the above directional GO pathways in the web-based SWAN platform.

#### SWAN Calculations

To simplify the discussion of the calculations, input annotations are referred to as genes and input data as copy-number alteration calls (−2 to 2 scale) for all genes for all tumors. In the first step, input data is subset for genes within the pathway of interest. As an example of what was applied to cancer data, this would be copy-number alteration calls for each tumor’s set of genes within the pathway. A network data frame is next generated using protein-protein interaction data from BioGRID (Stark et al., 2006) by default or by the user’s custom interaction data. An integer 1 is then assigned to each gene in the pathway. Non-interacting genes are included as additional rows. If interactions are not scored, this data frame is created without edges and would then contain the same number of rows as genes in the pathway. Custom prioritization scores are then multiplied onto this network data frame (eg if a user decided to artificially prioritize 3-fold *ACTB* due to phenotype data, all nodes within the network table containing *ACTB* would be multiplied by 3). If haploinsufficiency is scored, this custom scoring includes a 2x multiplier for all genes with yeast haploinsufficient or triploproficient homologs and a 3x multiplier for all genes with a mouse haploinsufficient homolog. A pair of network scoring matrices is then generated for interaction pairs (a bait matrix and a prey matrix) by multiplying the input data by each integer value column of the interaction data frame. This step incorporates input data into the network scoring matrices.

For each pathway, two separate hypotheses are tested: the pathway is suppressed or the pathway is elevated. Mathematically, this only comes into the calculations for network interaction pairs with an unclear effect. This is when one node contains a negative value (eg, from gene deletion) and the interacting node contains a positive value (eg, from gene gain). The hypothesis is that if the pathway overall is driven toward elevation, then the positive value is the relevant value (eg, it exerts a dominant effect within the pathway and the loss of an interactor does not abrogate its function). Similarly, if the pathway overall is driven toward suppression, then the negative value is the relevant value (eg, it exerts a dominant effect within the pathway as a potential rate limiting factor for interacting proteins). Thus, a suppression matrix and an elevation matrix are both created to test each hypothesis. The exact form of accomplishing this is complex in form to aid in 100-fold speed improvements in the R programming environment and can be viewed in the supplied code. Briefly, the suppression matrix is created by taking the matrix position minimums between the network scoring matrices and zero, whereas the elevation matrix is created by taking the matrix position maximums between the network scoring matrices and zero. The network scores for each tumor are then summed across rows to form a pair of hypothesis vectors of network shift scores per tumor.

The process is then repeated in two control, randomly selected background subsets with an equal number of genes as the input subset. This yields two randomly permuted background vectors per hypothesis, which are used in future calculations to estimate network topology and provide a noise estimate. For example, the experimental / observed network shifts must be greater in magnitude and frequency than the shifts found from simply shuffling genes and comparing one set of shuffled genes to another set of shuffled genes. If an experimental control dataset is uploaded, these data take the place of one of the shuffled controls.

Now that network scores per tumor have been calculated for the actual observed data and two random controls, the differences between these are calculated. SWAN has two method options for this comparison. One is to create a single residual per tumor between the pathway genes’ data and the randomized genes’ data and subtract the residual of the second randomized genes’ data to the first randomized genes’ data. This residual is the difference of the sums of network edge scores. In the second method, residuals are created for each pathway gene within each tumor, compared to the randomized backgrounds of an equivalent number of genes, for each tumor. The reason there are two methods is to aid in analyzing appropriately sized datasets. In large datasets (N ≥ 15, as an approximation for CNA data), SWAN only requires single-residual-per-tumor data to reach a verdict for pathway suppression or elevation. In small datasets common to pilot murine or other complex and expensive biological experiments, the residuals acquired from each gene are required for SWAN to reach a significant verdict in pathway sets.

Since SWAN was designed to handle noisy data, further random permutations are then iteratively performed. In the presented data for this study, 1,000 total permutations are performed, and each of these has two randomly shuffled gene choice to build the random comparison networks (thus, 2,000 random networks per pathway). This is recommended to be reduced to N = 100 permutations to achieve more timely analysis without substantial loss of accurate pathway shift calls (**Figure S2D**). Permutations used in this study yielded 1,000 residual pairs to be used to determine which hypothesis was most likely for the queried pathway: was this pathway suppressed by CNAs or elevated by CNAs? We adopted a parsed majority vote method (Bouziane et al., 2011) to stabilize these hypothesis calls for pathways of marginally significant shifts. Briefly, the direction of the pathway shift is determined for each experimental tumor compared to each of the randomizations in sets of randomizations. We determined 25 randomizations per majority vote improved the stability of calls. For example, in borderline cases wherein 12 iterations were called as suppressed and 13 were called as elevated, the majority vote for this set was then issued as elevated. Majority votes per set of 25 (40 sub-majority votes total in our 1,000 permutation analyses) were then summed to determine the best-fit hypothesis for the given pathway: either suppression or elevation.

Once the pathway is determined as most likely suppressed or most likely elevated by this majority vote method, the statistical test is performed. A Wilcoxon rank-sum test is applied with appropriate directionality to the vector of residuals representing the tumor network shifts (the N here is the same as the number of samples / tumors) to the vector of residuals representing the random network shifts (same N as control). If a paired control dataset was used as input data, the Wilcoxon rank-sum test used includes the pairing setting. If gene-level-p is selected, the residuals of each gene per sample are instead used. This increases N to N = (# samples) * (# genes within the pathway). While the gene-level-p somewhat biases significance toward large pathways, this may be chosen as the logical route to analysis for small N experiments and is absolutely required to achieve significant false-discovery-rate (FDR) values as a nominal rank-sum test bottoms out at ∼0.1 for N = 4 samples using typical CNA data. Finally, in a pan-pathway analysis, the Wilcoxon *P*-values are multiple hypothesis testing FDR corrected using the Benjamini-Hochberg method or a q-value generated using a Bonferroni correction, allowing for all levels of stringency depending on varied user requirements.

Conceptual depictions of these calculations can be found in the documentation and videos accompanying SWAN software. For further details to these complex calculations, please refer to the available annotated code hosted at GitHub (https://github.com/jrdelaney/SWAN).

### Outputs

Tabular outputs from SWAN include summary data and individual data per sample. **(1)** A pathway summary file is created with one row per pathway tested. Columns for this tab-separated spreadsheet include (a) name of input file, (b) name of pathway, (c) result (suppressed = “Haploinsufficient”, elevated = “Triploproficient”, neither = “No Selection”), (d) the average shift of network score in the experimental group relative to the control group, (e) a nominal p-value by Wilcoxon ranksum, (f) a Bonferroni-corrected q value of these p-values, (g) an FDR correction of the p-values (Benjamini-Hochberg correction), (h) the standard deviation of network shift scores, and (i) prioritized genes along with their corresponding gene-specific network shift scores. **(2)** A per-sample file depicting network shift scores relative to control per pathway, for use in clustering and other algorithms. **(3)** A gene-centric output file depicting each gene’s influence across all pathways tested, which is another way to prioritize biologically impactful genes.

### Quality Control of SWAN

The cancer types used in the TSG and OG quality control (QC) analysis were ACC, BLCA, BRCA, CESC, COAD, ESGA, GBM, HNSC, KICH, KIRC, LGG, LIHC, LUAD, LUSC, MESO, OV, PAAD, PRAD, READ, SARC, SKCM, STAD, TGCT, THCA, UCEC, and UVM.

To test whether SWAN was able to enrich for known oncogenes and tumor suppressor genes in its list of prioritized genes, tabular results from SWAN were queried. COSMIC Cancer Gene Census Tier 1 was used as the list of tested OGs and TSGs. As a possible alternative, STOP and GO genes were also tested. Each STOP and GO gene used was from two of three sources from the Elledge lab (Davoli et al., 2013; Sack et al., 2018; Solimini et al., 2012). For “gene prioritization enrichment,” genes from pathways called as “haploinsufficient” or “triploproficient” (FDR ≤ 0.0001) marked by SWAN as in the top five negatively or positively scoring genes, respectively, were tabulated. The enrichment ratio is calculated as the ratio of the sum of TSGs in the top five prioritized genes within a haploinsufficient pathway divided by the sum of those which are not known TSGs, over a similar ratio within neutral-called (“no selection”) pathways from SWAN. An equivalent calculation was performed for OGs. A fisher’s exact test was performed on these counts for each cancer type separately for TSGs and OGs (**Figure 2B** and **Figure S3**). To determine the loss of enrichment, pan-cancer SWAN analysis was performed with 1,000 iterations across these 26 QC cancer types while adjusting single parameters. QC analysis was then performed again, and the percent difference in enrichment from the null 1 ratio value was calculated.

To determine an appropriate default cutoff FDR value for SWAN, a tuning range of “0.2,0.1,0.05,0.04,0.03,0.02,0.01” and then 10 fold less down to “10^-50^” was used. All 26 QC cancer types were again tested for a TSG and OG enrichment ratio. In this case, the fisher’s exact test summed TSGs and OGs together and non-TSGs with non-OGs together, comparing pathways called as significant to pathways called as non-significant. This allowed for a single metric which balanced sensitivity and specificity; the statistical test would yield a larger *P*-value for lower N of significant pathways as well as when the neutral-called pathways began to have similar rates of OGs and TSGs as significantly-called pathways. Scrolling across this metric, an optimal FDR ≤ 0.0001 was determined for a default value (**Figure S1E**).

### Gene Set Enrichment Analysis

26 cancer types were analyzed using Gene Set Enrichment Analysis (GSEA) version 4.0.3. Integer normalized TCGA data was used identically as in SWAN and diploid data was set as control. KEGG Pathway gene set was used with 1000 permutations to the phenotype us the “on-the-fly” setting. For each pathway, the top five enriched genes and the bottom five downregulated genes were considered prioritized to enable comparison with SWAN. The number and enrichment of tumor suppressor genes and oncogenes were calculated identically as defined in SWAN Quality Control.

### BET Inhibitor Study Analysis

Data sets used in each analysis were attained from Gene Expression Omnibus (GEO). Raw data files were formatted using SWAN Data Groomer and transformed to log base 2. Each resulting experimental and control data files were input into either pan-pathway or single pathway SWAN. Pan-pathway analysis calculated 200 control permutations and had a significance threshold of 0.05. Pathways including less than 10 or more than 200 genes were omitted. Single pathway analysis calculated 100 control permutations and had a significance threshold of 0.001

### Identification of novel oncogenes and tumor suppressors

SWAN Interactome summary data from GO pathway analysis was used as a starting point to classify genes as general CNA-influenced OGs or TSGs. Z-scores of each gene within the interactome dataset were averaged for OGs in cancers in which the Z-score was positive and a similar calculation was performed on negative values for TSGs. The number of times a gene was displayed on a cancer Interactome priority plot was summed for each candidate OG and TSG. Alpha transparency values represent how much CNA influence originates from the gene itself (ie, the gene is deleted or amplified) or from the interacting genes (ie, its interactors are deleted or amplified) and were tabulated for each gene in each tumor type. Only OGs and TSGs which were detected through this method in ≥ 5 tumor types are reported in the figures and supplemental tables. COSMIC Tier 1 genes were marked as “known” OGs and TSGs as long as they were not characterized as fusion-only TSGs or OGs. All genes outside these criteria were considered “novel”, although we caution this does not capture the entirety of the cancer literature. Essential gene data were downloaded from a 324 cell line CRISPR-Cas9 screen study (Behan et al., 2019). OGs which were not hits within this screen were included in the **Figure S6A**. TSGs as identified by SWAN interactome analysis not considered “known” OGs were plotted in **Figure 4D** and summarized within **Table S3**.

### Mutation association analysis

Genes mutated in >3% of all TCGA tumors were analyzed for differential SWAN shift distributions. Subsets were taken from each tumor type by splitting mutant versus non-mutant tumors by each putative driver gene. A Wilcoxon ranksum test was performed on SWAN shifts in mutant tumors and compared to non-mutant tumors. If the mean SWAN shift of the mutant group was lower than the mean SWAN shift of the non-mutant group, the association was considered negative and conversely for positive. The final Supplemental Table 6 lists pathways reaching an FDR (by Benjamini-Hochberg correction of *P*-values) less than 0.05. It may be noted that most genes only yielded significant associations in limited cancer types due to the inadequate number of mutations in other tumor types.

### Machine-learning prognostic analysis

Patients with both overall survival data and SWAN shift data were analyzed for prognostic SWAN pathways. Patient data were first separated by SWAN pathway shifts, with the “low” and “high” groups separated by 1 standard-deviation (SD) centered at the median shift. To consider the possibility of a false positive, 101 training and test sets were created for Cox-proportional hazard (CoxPH) models which utilize SWAN shift data. Each training set consists of a random selection of 67% of the tumors, and the test set is the remaining 33% of tumors. The training set is used to build a CoxPH model based on overall survival and SWAN shift scores and produce a hazard ratio (HR). This model is then applied to the test set to predict risk scores using SWAN shifts. The upper quartile risk is compared to the lower quartile risk group by log-rank test (survdiff of Surv function in the R package “survival”) to yield a *P*-value for the test group. Potential false positive prognostic pathways were removed by (1) determining significance in the applied CoxPH model risk for each test set, and (2) determining CoxPH HR direction in each training set. The most prognostic pathways are labeled as those with >80% of randomizations as significant and in same HR direction and provided as **Table S7**. Kaplan-Meier curves and survival analysis on the entire cohort without machine-learned filtering is available using the Shiny SWAN Single-Pathway app online.

### Cell culture and biologic quality control

Cells were cultured at 37°C with 5% CO_2_. Established cell lines (OVCAR3, SKOV3, CAOV3, and 293T) were purchased from the American Type Culture Collection (ATCC) and validated by short tandem repeat (STR) profiling. Routine microscopic morphology tests were performed prior to each experiment. All cells were grown in RPMI-1640 media supplemented with antibiotics (penicillin, streptomycin), sodium pyruvate, and 10% FBS (Gibco). Lentiviral constructs for *PEX5* and *PEX19* were purchased from Genecopoeia. *PEX19* cDNA (NM_002857.3) was cloned into EX-G0621-Lv242 with a C-Avi-FLAG tag and puromycin resistance. *PEX5* (NM_001131023.1) was synthesized and cloned into the EX-Z6463-Lv157 vector with C-3xHA Neomycin resistance. Lentivirus was produced in 293T cells and filtered through a 33µm filter prior to transduction. Confirmation of cDNA insert was performed by Sanger sequencing using forward primer 5’ AGGCACTGGGCAGGTAAG 3’ and reverse primer 5’ CTGGAATAGCTCAGAGGC 3’ for Lv242 and forward primer 5’ GCGGTAGGCGTGTACGGT 3’ and reverse primer 5’ ATTGTGGATGAATACTGCC 3’ for Lv157. SKOV3 and OVCAR3 cells were selected for Lv242 lentiviral integration by addition of 4µg/mL puromycin (Thermo Fisher) to the media or for Lv157 integration by addition of 200µg/mL Geneticin (Fisher Scientific).

For determination of mRNA expression of metallothionein isoforms, 1×10^6^ CAOV3 or 2×10^6^ OVCAR3 cells per well were plated in a 6-well dish. After 20h, cells were rinsed once with PBS (phosphate-buffered saline), and RNA was isolated using the miRNeasy Mini Kit (Qiagen) according to the protocol of the manufacturer. One µg of total RNA was used to transcribe cDNA using the iScript cDNA Synthesis Kit (Biorad). For quantitative real-time reverse-transcriptase polymerase chain reaction (qPCR) 20ng of cDNA per reaction and the iTaq Universal SYBR Green Supermix (Biorad) was employed. Triplicate samples were normalized to TBP and relative gene expression was determined by the ΔΔCt method. To knock down metallothionein 2A (*MT2A*) mRNA expression, shRNA lentiviral vectors targeting MT2A and scrambled control were used to generate lentivirus in 293T cells and filtered through a 33µm filter prior to transduction. CAOV3 and OVCAR3 cells were selected for lentiviral integration by addition of 4µg/mL puromycin (Thermo Fisher). For γ H2AX staining 2.5×10^3^ CAOV3 or OVCAR3 cells were seeded onto black, optical bottom, 96-well plates (VWR). After allowing the cells to adhere for 16h, cells were treated with 50 µM (CAOV3) or 100 µM (OVCAR3) CdCl_2_ (Sigma-Aldrich) for 24h. Then cells were fixed in 4% paraformaldehyde for 10min, permeabilized with 0.1% Triton X-100 for 2min, and nonspecific binding was blocked with PBS containing 5% bovine serum albumin and 5% goat serum for 45min. Then cells were incubated with purified anti-γH2AX antibody (phospho-Ser139; BioLegend) overnight, primary antibody was removed with three PBS washes of 10 min, incubated in Hoechst 33342 (Fisher Scientific) and secondary goat anti-mouse Alexa Fluor 594nm antibody (LifeTechnologies) for 1.5h, and secondary was removed followed by three PBS washes of 10 min each. Finally, a Lionheart FX automated microscope (BioTek) was used to image the cells and ImageJ was employed to quantify γH2AX puncta number and intensity.

### Metabolomics

Metabolomics were performed as previously described (Delaney et al., 2020). Specific changes for the data presented here include: five replicates were used per genetic condition, cell line used was SKOV3. Otherwise, the methods are repeated and copied below.

All samples were grown to 80% confluency on a 10cm tissue culture dish. Cells were harvested by trypsinization and neutralized with iced RPMI complete media. Cells were washed twice in iced PBS and split into two tubes. Cell pellets were saved at −80°C until analysis. All sample sets had three independent cell growth experiments performed on different days.

For amino acid and lipid analysis, tubes containing ovarian cancer cell pellets were thawed at room temperature and then stored on ice during manipulation. For normalization, a duplicate pellet was analyzed by BCA assay for total content determination. 100 µL of 80/20 v/v MeOH/water was added to each sample tube. Samples were then probe sonicated 3 times at power level 3 for 5 seconds each burst, cooling on ice between bursts. Samples were then allowed to incubate for 10 minutes while on ice and then put in −80°C freezer until ready for analysis.

Samples were prepared using the AbsoluteIDQ p180 kit (Biocrates Innsbruck, Austria) in strict accordance with their detailed protocol. Samples were taken from the −80°C freezer and centrifuged at 4°C for 10 minutes at 15,000g. After the addition of 10 µL of the supplied internal standard solution to each well of the 96-well extraction plate, 15 µL of each ovarian study sample was added to the appropriate wells. The plate was then dried under a gentle stream of nitrogen for 10 minutes. An additional 15 µL of each study sample was added to the respective wells and plate was dried under nitrogen for an additional 20 minutes. The samples were derivatized with phenyl isothiocyanate then eluted with 5mM ammonium acetate in methanol. Samples were diluted with either 1:1 methanol:water for the UPLC analysis (4:1) or running solvent (a proprietary mixture provided by Biocrates) for flow injection analysis (20:1).

A study pool sample was created (5041 SPQC) by taking an equal volume from each study sample. The pooled sample was prepared and analyzed in the same way as the study samples in triplicate. On the kit plate, the SPQC was prepared in triplicate; one of these preparations was analyzed in triplicate while the other two were analyzed in a staggered manner before, during, and after the study samples in order to measure the performance of the assay across the sample cohort. The five analyses of this pool can be used to assess potential quantitative drift across the analysis of the plate, or in larger studies, to assess batch effects.

UPLC separation of amino acids and biogenic amines was performed using a Waters (Milford, MA) Acquity UPLC with a Waters Acquity 2.1 mm x 50 mm 1.7 µm BEH C18 column fitted with a Waters Acquity BEH C18 1.7 µm Vanguard guard column. Analytes were separated using a gradient from 0.2% formic acid in water, to 0.2% formic acid in acetonitrile. Total UPLC analysis time was approximately 7 minutes per sample. Acylcarnitines, sphingolipids, and glycerophospholipids were analyzed by flow injection analysis (FIA) with total analysis time of approximately 3 minutes per sample. Using electrospray ionization in positive mode, samples for both UPLC and flow injection analysis were introduced directly into a Xevo TQ-S triple quadrupole mass spectrometer (Waters) operating in the Multiple Reaction Monitoring (MRM) mode. MRM transitions (compound-specific precursor to product ion transitions) for each analyte and internal standard were collected over the appropriate retention time. The UPLC-MS/MS data were imported into Waters application TargetLynx for peak integration, calibration, and concentration calculations. The UPLC-MS/MS data from TargetLynx and FIA-MS/MS data were analyzed using Biocrates MetIDQ software. For statistical comparisons of glycerophospholipids and sphingolipids, a Wilcoxon rank-sum test was performed. All other tests were a student’s t-test.

For the energy metabolites including Acetyl-CoA, NAD+, Glutathione, cAMP, AMP, ADP, and ATP, an alternate assay was performed on the same cell pellet following the sonication step. The samples were then placed in a cold aluminum sample block on dry ice and incubated for 10 minutes. Next the samples were centrifuged for 10 minutes at 4°C and 15,000 g and stored at −80°C until ready for analysis. The samples were warmed to 4°C on ice and centrifuged again for 10 minutes at 4°C and 15,000 g to pellet any solids. Forty microliters of supernatant from each sample was pipetted into a glass total recovery vial (Waters) labeled with its corresponding DPMSR ID number. The remaining pellet from each sample was stored at −80°C. A study pool quality control (SPQC) sample was prepared by combining 5 µL of supernatant from each sample into a 1.5 mL tube (Eppendorf). Stable Isotope Labeled (SIL) standard material, the 13C Credentialed E. coli kit (MS-CRED-KIT) was purchased from Cambridge Isotope Laboratories. This is an E. coli extract from uniformly 13C-labeled E. coli. The material was tested to have minimal to no contributing signal in the light channel, using injections of only the 13C-labeled standard in previous experiments. Nine hundred microliters (900 µL) of sample resuspension solvent was created by taking one vial of 13C-labeled MS-CRED-KIT containing 100 µL of lysate and adding 400 µL of 80:20 v/v methanol/water. This resuspension solution was prepared immediately before addition to the samples. This solution was also used as the internal standard blank during the analysis. Ten microliters of the 13C-labeled *E. coli* resuspension solution was added via repeater pipette to each sample in the glass total recovery vials. Four microliters from each sample was injected for analysis by LC-MS/MS.

Liquid chromatographic separation was performed using a Waters Acquity UPLC with a 2.1 mm x 30 mm, 1.7 μm pore size ethylene bridged hybrid (BEH) amide column (Waters PN: 186004839). Mobile phase A was composed of water with 10 mM ammonium hydrogen carbonate (AmBic) (Millipore Sigma, St. Louis, MO) containing 0.2% ammonium hydroxide (NH_4_OH) generated as follows: 3.34 mL of 30% ACS grade NH_4_OH was added to 1 L water, followed by the addition of 0.3982 g AmBic. Mobile phase B was neat acetonitrile (Optima LCMS grade Thermo). The weak needle wash was mobile phase B and the strong needle wash was mobile phase A. The total length of the LC Gradient Program is 5.00 minutes. The outlet of the analytical column was connected directly via electrospray ionization into a Xevo TQ-S mass spectrometer (Waters) with positive/negative mode switching. Retention time scheduling with 30 second windows was used to minimize concurrent MRM transitions, and automatic dwell calculation was used to maximize dwell time while maintaining at least 8 points across the chromatographic peak. Eighty milliseconds (80 msec) was set as the polarity-switching delay. Positive and negative ion electrospray were alternated during the entirety of an LC gradient program for one injection. In ESI+ mode, capillary voltage was 3.0kV, source offset was 50V, desolvation temperature was 400°C, desolvation gas flow was 650 L/hr N2, cone gas was 150 L/hr N2, and nebulizer pressure of 7.0 bar was used. Source parameters for ESI-ionization were the same as ESI+, with the exception of the capillary voltage was set to −2.0 kV. Each sample was analyzed in Multiple Reaction Monitoring (MRM) mode in the mass spectrometer during the LC gradient program as ions eluted from the LC column. Metabolomic data associated with the figures is provided in **Table S4**.

### Flow cytometry

SKOV3 and OVCAR3 cells transduced with LV157 or LV242 with or without *PEX5* and *PEX19* respectively were seeded at 25,000 cells per well in a 24 well TC plate in 1mL media containing antibiotics. A day after seeding, cisplatin (10µM in DMSO) or N-acetyl-cysteine (2mM in ddH2O) were added to the media and the cells grown for 48h prior to staining for flow cytometry. Staining was performed with 10µM H_2_DCFDA (2′,7′-Dichlorodihydrofluorescein diacetate, VWR #89138-260) for 1 hour. Cells were then rinsed with PBS and 500µL Trypsin 0.05% EDTA (Thermo Fisher Scientific #25300120) was added for 5 minutes. Trypsinized cells were added to 500µL iced RPMI in 1.5mL microcentrifuge tubes. Cells were centrifuged at 3,000g for 1 minute and media aspirated. 1mL iced PBS was then added to cells and cells were briefly resuspended. Cells were centrifuged at 3,000g for 1 minute. PBS was aspirated, 300µL fresh iced PBS was added to cells and cells were transferred to an iced 5mL polypropylene flow cytometry tube (VWR #352063). Cells were analyzed for fluorescence in the 488nm channel on a BD FACSCanto II cytometer and analyzed using BD FACSDiva software.

### Cell death and proliferation assays

SKOV3 and OVCAR3 cells transduced with LV157 or LV242 with or without *PEX5* and *PEX19* respectively were seeded at 10,000 cells per well in a 96 well TC plate in 50µL media containing antibiotics. Cells were allowed to adhere for 3 hours prior to addition of 50µL media containing 2X treatment solution (20µM cisplatin, 4mM NAC, and/or 0.2% DMSO control). Cells were then grown for 48h prior to fixation. For fixation, cells were first rinsed in 125µL PBS and then stained with crystal violet staining and fixation solution (0.11% crystal violet, 0.17M NaCl, and 22% methanol in ddH2O) for 15 minutes. Crystal violet stain was aspirated, 125µL PBS wash performed twice, and then cells were dried for 30 minutes at 37°C in a dry incubator. 85µL methanol was added to each well and absorbance was read in an absorbance spectrophotometer at 600nm. Background consisting of cells killed to 100% penetrance using 1mM H_2_O_2_ was subtracted from all reads. Growth inhibition was calculated as the fractional difference in absorbance of a treated well compared to the average control-treated well for an isogenic cell line on the same 96-well plate.

### Whole-exome sequencing and data processing

Samples were processed using a Promega Maxwell RSC Instrument (AS4500) and Maxwell RSC Tissue DNA kit (AS1610) to obtain purified DNA. DNA was sent to GENEWIZ for whole-exome processing using an Agilent SureSelect Human All Exon V6 kit and next-generation sequencing on an Illumina HiSeq-4000.

SNVs and indels in *TP53* were called using one of two methods. The first method was a default DRAGEN protocol used by GENEWIZ. The second, used to call mutations in the remaining half of samples, utilized a triple-tool calling method. FASTQ reads were aligned to hg38 to create BAM files. BAM files were removed of PCR duplicated using RmDup. The three variant callers used on the BAM files were: LoFreq, FreeBayes, and samtools followed by VarScan (Koboldt et al., 2009; Wilm et al., 2012). Variant callers were run using the Galaxy platform (Afgan et al., 2016). *TP53* mutations called by all three tools were then filtered by those present in gnomAD v3 (Karczewski et al., 2020) at an allele frequency > 0.0001 in any ethnicity group. Annovar was used to annotate variants (Wang et al., 2010). To query ethnicity at genome-scale, EthSEQ (Romanel et al., 2017) was used on genome-wide called variants fitting the three-tool intersect threshold without a gnomAD cutoff (N = 7,695 variants assessed in new cohort).

ControlFREEC (Boeva et al., 2012) was used to call CNAs without using paired normal controls. To aid in estimation of stromal cell contamination, the TP53 mutation allele frequency was used as the estimated tumor cell fraction. Settings included: breakPointThreshold (1.2), readCountThreshold (50), window (500,000bp), telocentromeric (100,000bp), contaminationAdjustment (TRUE), sex (XX), contamination (using normal *TP53* allele fraction), and a single control uterus was used as a control target capture region. To best match TCGA CNA data, all BAMs used were aligned to hg19. To create a −2 to 2 normalized file, negative CNAs were given a value of −1, positive CNAs a value of +1, and any positive CNAs exceeding 2 standard deviations above the median a value of +2. Gene-level CNAs were determined using the SWAN Data Groomer functions built for *.seg files. Genome conversion of CNAs or mutation variants between hg19 and hg38 used liftOver with UCSC chain files (Kuhn et al., 2013).

### Quantification and Statistical Analysis

In all cell biology figures, *P*-values are calculated using a two-tailed Student’s t-test, unless otherwise indicated. The description of SWAN describes SWAN statistical considerations in detail. Survival outcomes were assessed using Kaplan-Meier curves with log-rank tests.

### Resource Availability

#### Lead Contact

Further information and requests for resources and reagents should be directed to and will be fulfilled by the Lead Contact, Joe Delaney, PhD

#### Materials Availability

Cell lines and plasmids generate in this study are maintained at the Medical University of South Carolina and will be made available upon request.

#### Data and Code Availability

Protected human datasets generated during this study are available in dbGaP, accession phs002313. The code generated during this study are available at Github (https://github.com/jrdelaney/SWAN) and web software at https://www.delaneyapps.com/#SWAN.

## Declarations

This project was supported by the South Carolina Clinical & Translational Research Institute with an academic home at the Medical University of South Carolina CTSA NIH NCATS grant numbers UL1TR001450 (JRD) and TL1TR001451 (JKB). This work was supported by NIH grants CA207729 (JRD), GM119512 (DTL), and GM132055 (CMJ). The MUSC Proteogenomics Facility was used and is supported by GM103499 and MUSC’s Office of the Vice President for Research. Supported in part by the Biostatistics Shared Resource, Hollings Cancer Center, Medical University of South Carolina (P30 CA138313). The funders had no role in study design, data collection and analysis, decision to publish, or preparation of the manuscript. The contents are solely the responsibility of the authors and do not necessarily represent the official views of the NIH, NCI, or NCATS.

## Author Contributions

J.R.D., E.A.P., D.T.L., and R.R.B. designed the research. K.E.A. advised statistical considerations. J.R.D., R.R.B., and C.M.J. wrote the paper. J.R.D. scripted SWAN. All authors performed the research and analyzed the data.

## Supporting information

STAR Key Resources Table

Comparison of GSEA and SWAN for prioritization of known oncogenes and tumor suppressor genes

Pan-cancer SWAN pathway analysis summary

Novel oncogenes and tumor suppressor CNA driver genes identified by SWAN interactome analysis

Metabolomic data of PEX19 overexpression

New African American ovarian cancer cohort data and primers used

Summary of CNA pathways significantly associated with driver SNVs

Summary of SWAN pathways passing machine-learning prognostic threshold

## References

1. Afgan, E., Baker, D., van den Beek, M., Blankenberg, D., Bouvier, D., Cech, M., Chilton, J., Clements, D., Coraor, N., Eberhard, C., et al. (2016). The Galaxy platform for accessible, reproducible and collaborative biomedical analyses: 2016 update. Nucleic Acids Res 44, W3–W10.

2. Bakhoum, S.F., Ngo, B., Laughney, A.M., Cavallo, J.A., Murphy, C.J., Ly, P., Shah, P., Sriram, R.K., Watkins, T.B.K., Taunk, N.K., et al. (2018). Chromosomal instability drives metastasis through a cytosolic DNA response. Nature 553, 467–472.

3. Behan, F.M., Iorio, F., Picco, G., Goncalves, E., Beaver, C.M., Migliardi, G., Santos, R., Rao, Y., Sassi, F., Pinnelli, M., et al. (2019). Prioritization of cancer therapeutic targets using CRISPR-Cas9 screens. Nature 568, 511–516.

4. Ben-David, U., and Amon, A. (2019). Context is everything: aneuploidy in cancer. Nat Rev Genet.

5. Beroukhim, R., Mermel, C.H., Porter, D., Wei, G., Raychaudhuri, S., Donovan, J., Barretina, J., Boehm, J.S., Dobson, J., Urashima, M., et al. (2010). The landscape of somatic copy-number alteration across human cancers. Nature 463, 899–905.

6. Boeva, V., Popova, T., Bleakley, K., Chiche, P., Cappo, J., Schleiermacher, G., Janoueix-Lerosey, I., Delattre, O., and Barillot, E. (2012). Control-FREEC: a tool for assessing copy number and allelic content using next-generation sequencing data. Bioinformatics 28, 423–425.

7. Bouziane, H., Messabih, B., and Chouarfia, A. (2011). Profiles and majority voting-based ensemble method for protein secondary structure prediction. Evol Bioinform Online 7, 171–189.

8. Bowry, A., Piberger, A.L., Rojas, P., Saponaro, M., and Petermann, E. (2018). BET Inhibition Induces HEXIM1- and RAD51-Dependent Conflicts between Transcription and Replication. Cell Rep 25, 2061–2069 e2064.

9. Bozic, I., Reiter, J.G., Allen, B., Antal, T., Chatterjee, K., Shah, P., Moon, Y.S., Yaqubie, A., Kelly, N., Le, D.T., et al. (2013). Evolutionary dynamics of cancer in response to targeted combination therapy. Elife 2, e00747.

10. Cai, Y., Crowther, J., Pastor, T., Abbasi Asbagh, L., Baietti, M.F., De Troyer, M., Vazquez, I., Talebi, A., Renzi, F., Dehairs, J., et al. (2016). Loss of Chromosome 8p Governs Tumor Progression and Drug Response by Altering Lipid Metabolism. Cancer Cell 29, 751–766.

11. Cancer Genome Atlas Research, N. (2011). Integrated genomic analyses of ovarian carcinoma. Nature 474, 609–615.

12. Cancer Genome Atlas Research, N., Brat, D.J., Verhaak, R.G., Aldape, K.D., Yung, W.K., Salama, S.R., Cooper, L.A., Rheinbay, E., Miller, C.R., Vitucci, M., et al. (2015). Comprehensive, Integrative Genomic Analysis of Diffuse Lower-Grade Gliomas. N Engl J Med 372, 2481–2498.

13. Castellano, E., and Downward, J. (2011). RAS Interaction with PI3K: More Than Just Another Effector Pathway. Genes Cancer 2, 261–274.

14. Crespo, I., Gotz, L., Liechti, R., Coukos, G., Doucey, M.A., and Xenarios, I. (2016). Identifying biological mechanisms for favorable cancer prognosis using non-hypothesis-driven iterative survival analysis. NPJ Syst Biol Appl 2, 16037.

15. Davoli, T., Xu, A.W., Mengwasser, K.E., Sack, L.M., Yoon, J.C., Park, P.J., and Elledge, S.J. (2013). Cumulative haploinsufficiency and triplosensitivity drive aneuploidy patterns and shape the cancer genome. Cell 155, 948–962.

16. Delaney, J.R., Patel, C.B., Bapat, J., Jones, C.M., Ramos-Zapatero, M., Ortell, K.K., Tanios, R., Haghighiabyaneh, M., Axelrod, J., DeStefano, J.W., et al. (2020). Autophagy gene haploinsufficiency drives chromosome instability, increases migration, and promotes early ovarian tumors. PLoS Genet 16, e1008558.

17. Delaney, J.R., Patel, C.B., Willis, K.M., Haghighiabyaneh, M., Axelrod, J., Tancioni, I., Lu, D., Bapat, J., Young, S., Cadassou, O., et al. (2017). Haploinsufficiency networks identify targetable patterns of allelic deficiency in low mutation ovarian cancer. Nat Commun 8, 14423.

18. Du, Z., Song, X., Yan, F., Wang, J., Zhao, Y., and Liu, S. (2018). Genome-wide transcriptional analysis of BRD4-regulated genes and pathways in human glioma U251 cells. Int J Oncol 52, 1415–1426.

19. Figueiredo, C.R., Kalirai, H., Sacco, J.J., Azevedo, R.A., Duckworth, A., Slupsky, J.R., Coulson, J.M., and Coupland, S.E. (2020). Loss of BAP1 expression is associated with an immunosuppressive microenvironment in uveal melanoma, with implications for immunotherapy development. J Pathol 250, 420–439.

20. Flaherty, K.T., Gray, R., Chen, A., Li, S., Patton, D., Hamilton, S.R., Williams, P.M., Mitchell, E.P., Iafrate, A.J., Sklar, J., et al. (2020). The Molecular Analysis for Therapy Choice (NCI-MATCH) Trial: Lessons for Genomic Trial Design. J Natl Cancer Inst 112, 1021–1029.

21. Gambacorti-Passerini, C. (2008). Part I: Milestones in personalised medicine--imatinib. Lancet Oncol 9, 600.

22. Gao, J., Aksoy, B.A., Dogrusoz, U., Dresdner, G., Gross, B., Sumer, S.O., Sun, Y., Jacobsen, A., Sinha, R., Larsson, E., et al. (2013). Integrative analysis of complex cancer genomics and clinical profiles using the cBioPortal. Sci Signal 6, pl1.

23. Garcia-Carpizo, V., Ruiz-Llorente, S., Sarmentero, J., Grana-Castro, O., Pisano, D.G., and Barrero, M.J. (2018). CREBBP/EP300 bromodomains are critical to sustain the GATA1/MYC regulatory axis in proliferation. Epigenetics Chromatin 11, 30.

24. Group, P.T.C., Calabrese, C., Davidson, N.R., Demircioglu, D., Fonseca, N.A., He, Y., Kahles, A., Lehmann, K.V., Liu, F., Shiraishi, Y., et al. (2020). Genomic basis for RNA alterations in cancer. Nature 578, 129–136.

25. Hildebrand, J.S., Wallace, K., Graybill, W.S., and Kelemen, L.E. (2019). Racial disparities in treatment and survival from ovarian cancer. Cancer Epidemiol 58, 77–82.

26. Kang, M.S., Kim, J., Ryu, E., Ha, N.Y., Hwang, S., Kim, B.G., Ra, J.S., Kim, Y.J., Hwang, J.M., Myung, K., et al. (2019). PCNA Unloading Is Negatively Regulated by BET Proteins. Cell Rep 29, 4632–4645 e4635.

27. Karczewski, K.J., Francioli, L.C., Tiao, G., Cummings, B.B., Alfoldi, J., Wang, Q., Collins, R.L., Laricchia, K.M., Ganna, A., Birnbaum, D.P., et al. (2020). The mutational constraint spectrum quantified from variation in 141,456 humans. Nature 581, 434–443.

28. Kim, P.K., and Hettema, E.H. (2015). Multiple pathways for protein transport to peroxisomes. J Mol Biol 427, 1176–1190.

29. Klaassen, C.D., Liu, J., and Choudhuri, S. (1999). Metallothionein: an intracellular protein to protect against cadmium toxicity. Annu Rev Pharmacol Toxicol 39, 267-294.

30. Koboldt, D.C., Chen, K., Wylie, T., Larson, D.E., McLellan, M.D., Mardis, E.R., Weinstock, G.M., Wilson, R.K., and Ding, L. (2009). VarScan: variant detection in massively parallel sequencing of individual and pooled samples. Bioinformatics 25, 2283–2285.

31. Kuhn, R.M., Haussler, D., and Kent, W.J. (2013). The UCSC genome browser and associated tools. Brief Bioinform 14, 144–161.

32. Kumar, M., Bowers, R.R., and Delaney, J.R. (2020a). Single-cell analysis of copy-number alterations in serous ovarian cancer reveals substantial heterogeneity in both low- and high-grade tumors. Cell Cycle, 1-13.

33. Kumar, S., Warrell, J., Li, S., McGillivray, P.D., Meyerson, W., Salichos, L., Harmanci, A., Martinez-Fundichely, A., Chan, C.W.Y., Nielsen, M.M., et al. (2020b). Passenger Mutations in More Than 2,500 Cancer Genomes: Overall Molecular Functional Impact and Consequences. Cell 180, 915–927 e916.

34. Lee-Six, H., Olafsson, S., Ellis, P., Osborne, R.J., Sanders, M.A., Moore, L., Georgakopoulos, N., Torrente, F., Noorani, A., Goddard, M., et al. (2019). The landscape of somatic mutation in normal colorectal epithelial cells. Nature 574, 532–537.

35. Leone, R.D., and Emens, L.A. (2018). Targeting adenosine for cancer immunotherapy. J Immunother Cancer 6, 57.

36. Liberzon, A., Subramanian, A., Pinchback, R., Thorvaldsdottir, H., Tamayo, P., and Mesirov, J.P. (2011). Molecular signatures database (MSigDB) 3.0. Bioinformatics 27, 1739–1740.

37. Liu, Y., Chen, C., Xu, Z., Scuoppo, C., Rillahan, C.D., Gao, J., Spitzer, B., Bosbach, B., Kastenhuber, E.R., Baslan, T., et al. (2016). Deletions linked to TP53 loss drive cancer through p53-independent mechanisms. Nature 531, 471–475.

38. Long, Y., Sou, W.H., Yung, K.W.Y., Liu, H., Wan, S.W.C., Li, Q., Zeng, C., Law, C.O.K., Chan, G.H.C., Lau, T.C.K., et al. (2019). Distinct mechanisms govern the phosphorylation of different SR protein splicing factors. J Biol Chem 294, 1312–1327.

39. Longo, D.L. (2019). Personalized Medicine for Primary Treatment of Serous Ovarian Cancer. N Engl J Med 381, 2471–2474.

40. Ma, F., Li, B., Liu, S.Y., Iyer, S.S., Yu, Y., Wu, A., and Cheng, G. (2015). Positive feedback regulation of type I IFN production by the IFN-inducible DNA sensor cGAS. J Immunol 194, 1545–1554.

41. Mamlouk, S., Childs, L.H., Aust, D., Heim, D., Melching, F., Oliveira, C., Wolf, T., Durek, P., Schumacher, D., Blaker, H., et al. (2017). DNA copy number changes define spatial patterns of heterogeneity in colorectal cancer. Nat Commun 8, 14093.

42. Mannick, J.B., Del Giudice, G., Lattanzi, M., Valiante, N.M., Praestgaard, J., Huang, B., Lonetto, M.A., Maecker, H.T., Kovarik, J., Carson, S., et al. (2014). mTOR inhibition improves immune function in the elderly. Sci Transl Med 6, 268ra179.

43. Mannick, J.B., Morris, M., Hockey, H.P., Roma, G., Beibel, M., Kulmatycki, K., Watkins, M., Shavlakadze, T., Zhou, W., Quinn, D., et al. (2018). TORC1 inhibition enhances immune function and reduces infections in the elderly. Sci Transl Med 10.

44. Martincorena, I., Fowler, J.C., Wabik, A., Lawson, A.R.J., Abascal, F., Hall, M.W.J., Cagan, A., Murai, K., Mahbubani, K., Stratton, M.R., et al. (2018). Somatic mutant clones colonize the human esophagus with age. Science 362, 911–917.

45. Martincorena, I., Roshan, A., Gerstung, M., Ellis, P., Van Loo, P., McLaren, S., Wedge, D.C., Fullam, A., Alexandrov, L.B., Tubio, J.M., et al. (2015). Tumor evolution. High burden and pervasive positive selection of somatic mutations in normal human skin. Science 348, 880–886.

46. McElroy, J.A., Kruse, R.L., Guthrie, J., Gangnon, R.E., and Robertson, J.D. (2017). Cadmium exposure and endometrial cancer risk: A large midwestern U.S. population-based case-control study. PLoS One 12, e0179360.

47. Mitchell, T.J., Turajlic, S., Rowan, A., Nicol, D., Farmery, J.H.R., O’Brien, T., Martincorena, I., Tarpey, P., Angelopoulos, N., Yates, L.R., et al. (2018). Timing the Landmark Events in the Evolution of Clear Cell Renal Cell Cancer: TRACERx Renal. Cell 173, 611–623 e617.

48. Moore, L., Leongamornlert, D., Coorens, T.H.H., Sanders, M.A., Ellis, P., Dentro, S.C., Dawson, K.J., Butler, T., Rahbari, R., Mitchell, T.J., et al. (2020). The mutational landscape of normal human endometrial epithelium. Nature 580, 640–646.

49. Motohara, T., Masuda, K., Morotti, M., Zheng, Y., El-Sahhar, S., Chong, K.Y., Wietek, N., Alsaadi, A., Karaminejadranjbar, M., Hu, Z., et al. (2019). An evolving story of the metastatic voyage of ovarian cancer cells: cellular and molecular orchestration of the adipose-rich metastatic microenvironment. Oncogene 38, 2885–2898.

50. Nagarajan, S., Bedi, U., Budida, A., Hamdan, F.H., Mishra, V.K., Najafova, Z., Xie, W., Alawi, M., Indenbirken, D., Knapp, S., et al. (2017). BRD4 promotes p63 and GRHL3 expression downstream of FOXO in mammary epithelial cells. Nucleic Acids Res 45, 3130–3145.

51. Ohta, A., Gorelik, E., Prasad, S.J., Ronchese, F., Lukashev, D., Wong, M.K., Huang, X., Caldwell, S., Liu, K., Smith, P., et al. (2006). A2A adenosine receptor protects tumors from antitumor T cells. Proc Natl Acad Sci U S A 103, 13132–13137.

52. Onken, M.D., Li, J., and Cooper, J.A. (2014). Uveal melanoma cells utilize a novel route for transendothelial migration. PLoS One 9, e115472.

53. Patch, A.M., Christie, E.L., Etemadmoghadam, D., Garsed, D.W., George, J., Fereday, S., Nones, K., Cowin, P., Alsop, K., Bailey, P.J., et al. (2015). Whole-genome characterization of chemoresistant ovarian cancer. Nature 521, 489–494.

54. Poirier, Y., Antonenkov, V.D., Glumoff, T., and Hiltunen, J.K. (2006). Peroxisomal beta-oxidation--a metabolic pathway with multiple functions. Biochim Biophys Acta 1763, 1413–1426.

55. Prasad, V. (2020). Our best weapons against cancer are not magic bullets. Nature 577, 451.

56. Ren, C., Zhang, G., Han, F., Fu, S., Cao, Y., Zhang, F., Zhang, Q., Meslamani, J., Xu, Y., Ji, D., et al. (2018). Spatially constrained tandem bromodomain inhibition bolsters sustained repression of BRD4 transcriptional activity for TNBC cell growth. Proc Natl Acad Sci U S A 115, 7949–7954.

57. Romanel, A., Zhang, T., Elemento, O., and Demichelis, F. (2017). EthSEQ: ethnicity annotation from whole exome sequencing data. Bioinformatics 33, 2402–2404.

58. Rutledge, S.D., Douglas, T.A., Nicholson, J.M., Vila-Casadesus, M., Kantzler, C.L., Wangsa, D., Barroso-Vilares, M., Kale, S.D., Logarinho, E., and Cimini, D. (2016). Selective advantage of trisomic human cells cultured in non-standard conditions. Sci Rep 6, 22828.

59. Sack, L.M., Davoli, T., Li, M.Z., Li, Y., Xu, Q., Naxerova, K., Wooten, E.C., Bernardi, R.J., Martin, T.D., Chen, T., et al. (2018). Profound Tissue Specificity in Proliferation Control Underlies Cancer Drivers and Aneuploidy Patterns. Cell 173, 499–514 e423.

60. Santaguida, S., Vasile, E., White, E., and Amon, A. (2015). Aneuploidy-induced cellular stresses limit autophagic degradation. Genes Dev 29, 2010–2021.

61. Schrader, M., and Fahimi, H.D. (2006). Peroxisomes and oxidative stress. Biochim Biophys Acta 1763, 1755–1766.

62. Sheltzer, J.M., Ko, J.H., Replogle, J.M., Habibe Burgos, N.C., Chung, E.S., Meehl, C.M., Sayles, N.M., Passerini, V., Storchova, Z., and Amon, A. (2017). Single-chromosome Gains Commonly Function as Tumor Suppressors. Cancer Cell 31, 240–255.

63. Shi, Y., Ping, Y.F., Zhou, W., He, Z.C., Chen, C., Bian, B.S., Zhang, L., Chen, L., Lan, X., Zhang, X.C., et al. (2017). Tumour-associated macrophages secrete pleiotrophin to promote PTPRZ1 signalling in glioblastoma stem cells for tumour growth. Nat Commun 8, 15080.

64. Smith, J.C., and Sheltzer, J.M. (2018). Systematic identification of mutations and copy number alterations associated with cancer patient prognosis. Elife 7.

65. Solimini, N.L., Xu, Q., Mermel, C.H., Liang, A.C., Schlabach, M.R., Luo, J., Burrows, A.E., Anselmo, A.N., Bredemeyer, A.L., Li, M.Z., et al. (2012). Recurrent hemizygous deletions in cancers may optimize proliferative potential. Science 337, 104–109.

66. Sondka, Z., Bamford, S., Cole, C.G., Ward, S.A., Dunham, I., and Forbes, S.A. (2018). The COSMIC Cancer Gene Census: describing genetic dysfunction across all human cancers. Nat Rev Cancer 18, 696–705.

67. Song, X., Wan, X., Huang, T., Zeng, C., Sastry, N., Wu, B., James, C.D., Horbinski, C., Nakano, I., Zhang, W., et al. (2019). SRSF3-Regulated RNA Alternative Splicing Promotes Glioblastoma Tumorigenicity by Affecting Multiple Cellular Processes. Cancer Res 79, 5288–5301.

68. Stark, C., Breitkreutz, B.J., Reguly, T., Boucher, L., Breitkreutz, A., and Tyers, M. (2006). BioGRID: a general repository for interaction datasets. Nucleic Acids Res 34, D535–539.

69. Subramanian, A., Tamayo, P., Mootha, V.K., Mukherjee, S., Ebert, B.L., Gillette, M.A., Paulovich, A., Pomeroy, S.L., Golub, T.R., Lander, E.S., et al. (2005). Gene set enrichment analysis: a knowledge-based approach for interpreting genome-wide expression profiles. Proc Natl Acad Sci U S A 102, 15545–15550.

70. Sulzmaier, F.J., Jean, C., and Schlaepfer, D.D. (2014). FAK in cancer: mechanistic findings and clinical applications. Nat Rev Cancer 14, 598–610.

71. Suzuki, K., Kim, J.D., Ugai, K., Matsuda, S., Mikami, H., Yoshioka, K., Ikari, J., Hatano, M., Fukamizu, A., Tatsumi, K., et al. (2020). Transcriptomic changes involved in the dedifferentiation of myofibroblasts derived from the lung of a patient with idiopathic pulmonary fibrosis. Mol Med Rep 22, 1518–1526.

72. Taylor, A.M., Shih, J., Ha, G., Gao, G.F., Zhang, X., Berger, A.C., Schumacher, S.E., Wang, C., Hu, H., Liu, J., et al. (2018). Genomic and Functional Approaches to Understanding Cancer Aneuploidy. Cancer Cell 33, 676–689 e673.

73. Torres, E.M., Springer, M., and Amon, A. (2016). No current evidence for widespread dosage compensation in S. cerevisiae. Elife 5, e10996.

74. Verkman, A.S. (2011). Aquaporins at a glance. J Cell Sci 124, 2107–2112.

75. Wang, K., Li, M., and Hakonarson, H. (2010). ANNOVAR: functional annotation of genetic variants from high-throughput sequencing data. Nucleic Acids Res 38, e164.

76. Wedge, D.C., Gundem, G., Mitchell, T., Woodcock, D.J., Martincorena, I., Ghori, M., Zamora, J., Butler, A., Whitaker, H., Kote-Jarai, Z., et al. (2018). Sequencing of prostate cancers identifies new cancer genes, routes of progression and drug targets. Nat Genet 50, 682–692.

77. Wilm, A., Aw, P.P., Bertrand, D., Yeo, G.H., Ong, S.H., Wong, C.H., Khor, C.C., Petric, R., Hibberd, M.L., and Nagarajan, N. (2012). LoFreq: a sequence-quality aware, ultra-sensitive variant caller for uncovering cell-population heterogeneity from high-throughput sequencing datasets. Nucleic Acids Res 40, 11189–11201.

78. Yoshida, K., Gowers, K.H.C., Lee-Six, H., Chandrasekharan, D.P., Coorens, T., Maughan, E.F., Beal, K., Menzies, A., Millar, F.R., Anderson, E., et al. (2020). Tobacco smoking and somatic mutations in human bronchial epithelium. Nature 578, 266–272.

79. Zack, T.I., Schumacher, S.E., Carter, S.L., Cherniack, A.D., Saksena, G., Tabak, B., Lawrence, M.S., Zhang, C.Z., Wala, J., Mermel, C.H., et al. (2013). Pan-cancer patterns of somatic copy number alteration. Nat Genet 45, 1134–1140.

80. Zevini, A., Olagnier, D., and Hiscott, J. (2017). Crosstalk between Cytoplasmic RIG-I and STING Sensing Pathways. Trends Immunol 38, 194–205.

81. Zhang, H., Liu, T., Zhang, Z., Payne, S.H., Zhang, B., McDermott, J.E., Zhou, J.Y., Petyuk, V.A., Chen, L., Ray, D., et al. (2016). Integrated Proteogenomic Characterization of Human High-Grade Serous Ovarian Cancer. Cell 166, 755–765.

82. Zhang, J., Dulak, A.M., Hattersley, M.M., Willis, B.S., Nikkila, J., Wang, A., Lau, A., Reimer, C., Zinda, M., Fawell, S.E., et al. (2018). BRD4 facilitates replication stress-induced DNA damage response. Oncogene 37, 3763–3777.

