## Supplementary material for "SWAN Identification of Common Aneuploidy-Based Oncogenic Drivers": STAR Key Resources Table

| REAGENT or RESOURCE | SOURCE | IDENTIFIER |
| --- | --- | --- |
| Antibodies | | |
| H2A.X Phospho (Ser 139) | Biolegend | 613402 |
| Goat anti-Mouse IgG (H+L) Highly Cross-Adsorbed Secondary Antibody, Alexa Fluor 594 | Fisher Scientific | A-11032 |
| Bacterial and virus strains | | |
| DH5alpha | VWR | 101417-898 |
| NEB Stable Competent E. coli | NEB | C3040H |
| Biological samples | | |
| See Table S5, for flash frozen high-grade serous ovarian cancer tumor de-identified samples | Fox Chase Cancer Center, Cooperative Human Tissue Network | N/A |
| Chemicals, peptides, and recombinant proteins | | |
| acetic acid | Sigma-Aldrich | A6283 |
| bovine serum albumin | VWR | 97061-416 |
| cadmium chloride | Sigma-Aldrich | 202908 |
| cisplatin | Sigma-Aldrich | 232120 |
| dimethyl sulfoxide | Sigma-Aldrich | D8418 |
| EDTA | Fisher Scientific | 25300120 |
| Fetal Bovine Serum | Thermo Fisher | 10437028 |
| geneticin | Fisher Scientific | 10131035 |
| goat serum | VWR | 102643-594 |
| H2-DCFDA | VWR | 89138-260 |
| hoechst 33342, trihydrochloride, trihydrate | Fisher Scientific | H3570 |
| N-acetylcysteine | Sigma-Aldrich | 106425 |
| neomycin | VWR | 100219-896 |
| paraformaldehyde | Electron Microscopy Services | 15714 |
| Penicillin-Streptomycin | Sigma-Aldrich | P4333 |
| phosphate-buffered saline without Ca^++^ and Mg^++^ | VWR | 45000-446 |
| puromycin | Thermo Fisher | AAJ61278MB |
| RPMI 1640 with glutamine | VWR | 95042-508 |
| sodium pyruvate | Sigma-Aldrich | S8636-100ML |
| Triton X-100 | Sigma-Aldrich | X-100 |
| Critical commercial assays | | |
| AbsoluteIDQ kit | Biocrates | p180 |
| BCA Protein Assay Kit | Thermo Scientific | 23227 |
| iScript cDNA synthesis kit | Biorad | 1708891 |
| iTaq Universal SYBR Green Supermix | Biorad | 1725125 |
| Maxwell RSC Tissue DNA Kit | Promega | AS1610 |
| miRNeasy Mini Kit | Qiagen | 217004 |
| Optical Bottom Tissue Culture Plates, Greiner Bio-One | VWR | 82050-748 |
| ZymoPURE Plasmid Miniprep Kit | Zymo Research | D4209 |
| Deposited data | | |
| DNA sequencing assessment of CNVs of high-grade serous ovarian cancers | Database of Genotypes and Phenotypes (dbGaP) | dbGaP: phs002313 |
| Experimental models: cell lines | | |
| 293T | ATCC | CRL-3216 |
| CAOV3 | ATCC | HTB-75 |
| OVCAR3 | ATCC | HTB-161 |
| SKOV3 | ATCC | HTB-77 |
| Oligonucleotides | | |
| See **Table S5**, Primers tab | IDT | N/A |
| Recombinant DNA | | |
| pLv242-PEX19-OE (NM_002857.3) | Genecopoeia | EX-G0621-Lv242 |
| pLv157-PEX5-OE (NM_001131023.1) | Genecopoeia | EX-Z6463-Lv157 |
| pLv242-GFP (Vector 1) | Genecopoeia | pReceiver-Lv242 |
| pLv157-GFP (Vector 2) | Genecopoeia | pReceiver-Lv157 |
| pLKO.1 shMT2A-1 | Sigma | TRCN0000148975 |
| pLKO.1 shMT2A-2 | Sigma | TRCN0000148783 |
| TRC1/1.5 pLKO.1-puro Non-Target shRNA Control (shScr) | Sigma | SHC002 |
| Software and algorithms | | |
| R | R Project for Statistical Computing | https://www.r-project.org/  RRID:SCR_001905 |
| RStudio | RStudio | https://rstudio.com/  RRID:SCR_000432 |
| CAIRN | Delaney Lab | https://delaney.shinyapps.io/CAIRN/  RRID:SCR_019101 |
| FairSubset | Delaney Lab | https://delaney.shinyapps.io/FairSubset/  RRID:SCR_019102 |
| ImageJ software | http://rsbweb.nih.gov/ij/download.html | RRID:SCR_003070 |
