## Supplementary Figures and Legends for "SWAN Identification of Common Aneuploidy-Based Oncogenic Drivers"

### Supplemental Figure 1: SWAN concepts and optimizations

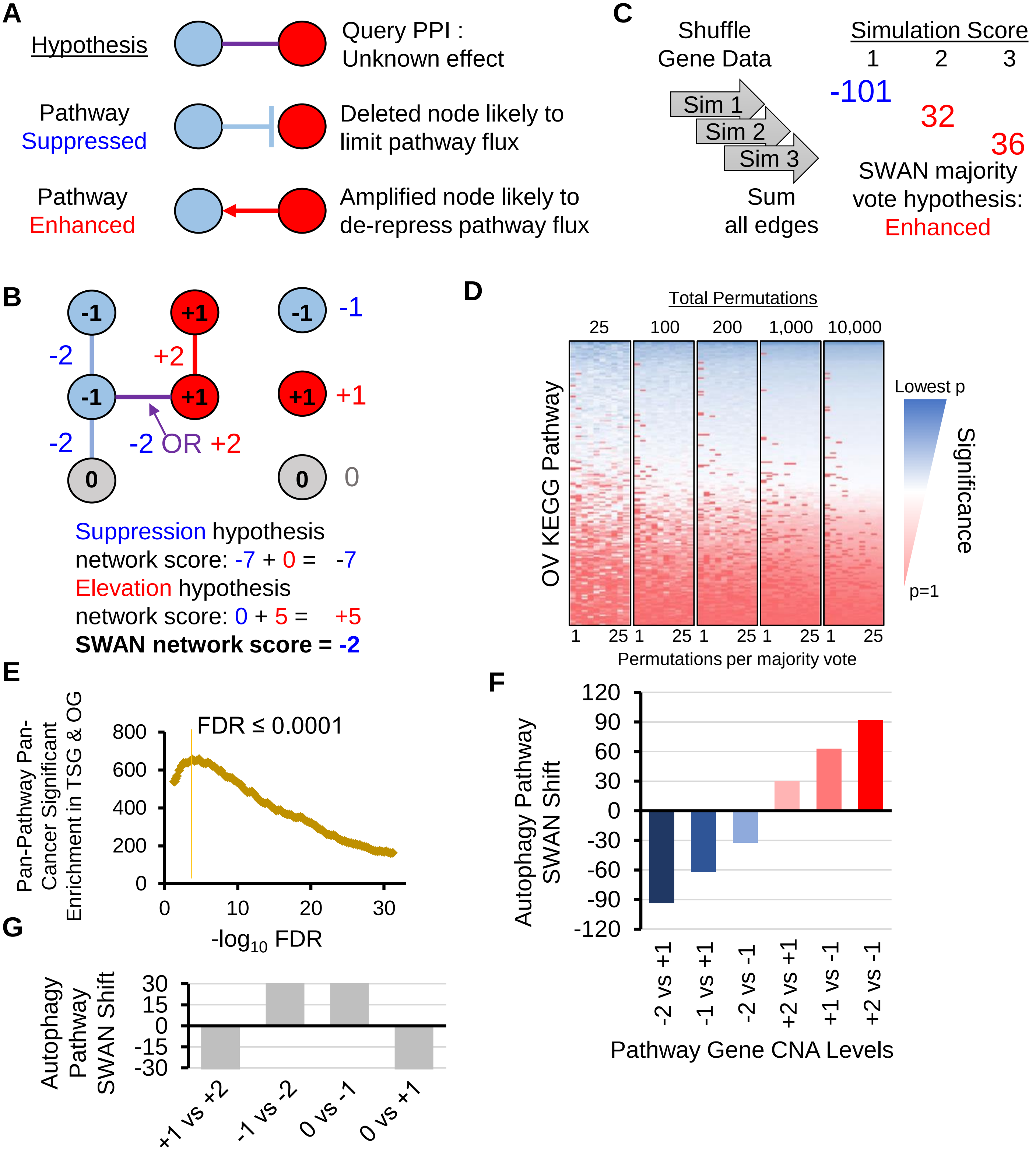

**Supplementary Figure 1. SWAN concepts and optimizations.** (**A**) Depiction of the two hypotheses that SWAN tests for each pathway with protein-protein interactions (PPI). The top scenario is the unknown: if one node is deleted (blue) and one node is amplified (red), how should the interaction be counted? If the pathway overall is selected for suppression, then the deleted node dominates by limiting pathway flux regardless of the amplified node (middle case). If the pathway overall is selected for elevation, then the amplified node dominates by circumventing the deletion (bottom case). (**B**) Mathematical calculations from the examples in (A) in the context of a simple pathway network. Grey indicates no CNA. Edges depict PPIs and nodes are genes. For each hypothesis, a SWAN network shift is calculated and shown here. Note that nodes without interactions are still scored, but with half the magnitude of a two-node edge. (**C**) An example of a simplified SWAN calculation using only 3 randomized background simulations (the default is 200, this work uses 1,000). By comparing the actual network (something similar to (B)) to networks generated using randomized background node data, SWAN shifts are generated. In this example, one shift was negative and two were positive, using the average of the two hypotheses for each summary score. In the majority vote calculation, the majority of SWAN shifts were positive, so an elevation hypothesis is used as the statistical test. Note that this avoids over-emphasizing extreme shifts, which would be the case here if the mean value was used, and stabilizes repeatable SWAN shift calls. (**D**) A graphical example of increased stability by majority voting, rather than increased permutations. With skewed network structures, some KEGG pathways using OV node data yield false negative SWAN $P$ = 1 without majority voting, even with 10,000 data randomizations for background estimates. However, with majority voting correction in sets of 25 randomizations, these skewed networks have very similar final SWAN P-values using only 100 permutations. (**E**) Graphical depiction of the sum of tumor suppressor gene (TSG) and oncogene (OG) enrichment values across all cancers tested, with suggested FDR cutoff value shown. (**F**) Simulated CNA data was generated for the KEGG: Autophagy pathway. A set the size of OV used OV CNA data, wherein autophagy gene CNAs were replaced by 50% “0” (no CNA) values and 50% x-axis indicated CNA value. A control set was identically created, using the x-axis “vs” numeric. SWAN shifts are stepped at a consistent magnitude, regardless of the nominal values, since the difference between experimental and control groups is a consistent shift. (**G**) A cautionary note on a common mis-interpretation of SWAN shifts using control data. Simulated pathway scores were generated using half of all genes with the CNA values shown on the x-axis. If a monoallelic deletion tumor set is compared to a homozygous deletion tumor set, then the monoallelic deletion tumor set will have a positive SWAN shift. Similar potentially counter-intuitive situations are shown. In all cases, it is recommended to view the CNA calls of SWAN prioritized genes to determine what genotype determined the calculated shift.

### Supplemental Figure 2: Quality Control of SWAN

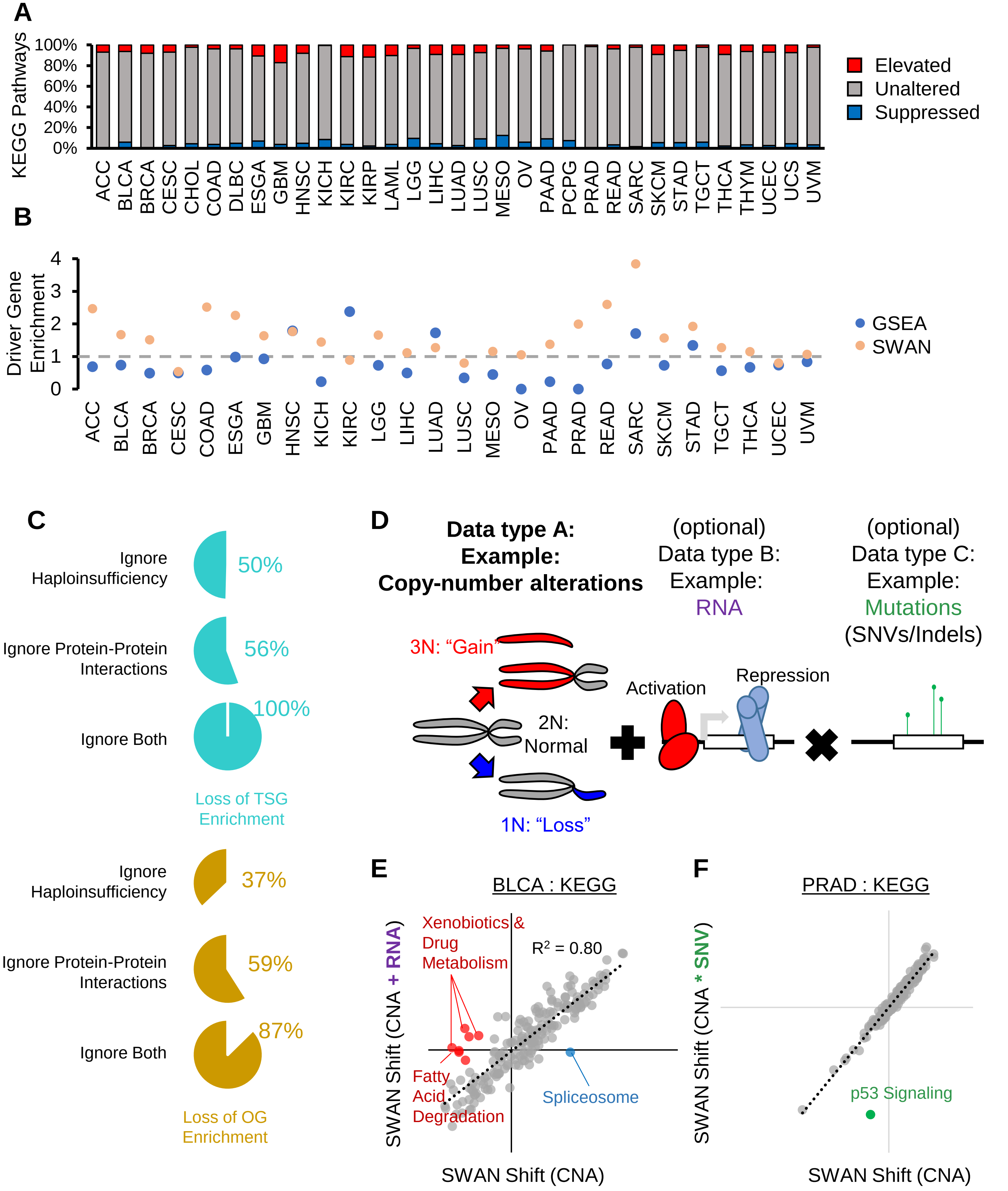

**Supplementary Figure 2. Quality control of SWAN.** Gene Set Enrichment Analysis of copy number alterations in 26 different cancer types, using KEGG pathway set. (**A**) Distribution of significantly ($p<0.05$) downregulated or upregulated pathways compared to not significantly ($p>0.05$) altered pathways. (**B**) Driver gene enrichment (mean of oncogene enrichment and tumor suppressor gene enrichment) in significantly altered pathways. See methods for details. An enrichment of >1 indicates enrichment of tumor suppressor genes or oncogenes in the significantly altered pathways relative to non-altered pathways. An enrichment of <1 indicates driver gene enrichment in unaltered pathways. An enrichment of 0 indicates no tumor suppressor genes or oncogenes were in the top five genes within the significantly altered pathways. (**C**) Pan-cancer SWAN tests were run without phenotypic data to determine if the phenotypes improve identification of OGs on elevated pathways and TSGs on suppressed pathways. Percentages represent the loss of >1 enrichment ratios relative to a null 1 value, averaged across all tumor types. (**D**) Conceptual example of how data layering (via additive or multiplicative layering) may be utilized in integrative SWAN pathway analysis. (**E**) An example of integrative RNA data layering with CNA data. Addition of similarly scaled RNA SWAN shifts to CNA data is plotted against SWAN shifts solely derived from CNA data for bladder cancer KEGG pathways. (**F**) An example of integrative mutation data layering with prostate adenocarcinoma CNA data. Mutations were multiplied into SWAN CNA networks and plotted against shifts generated solely from CNA data. The outlier p53 signaling pathway is highlighted.

### Supplemental Figure 3: Contribution of Phenotypes to SWAN Analysis

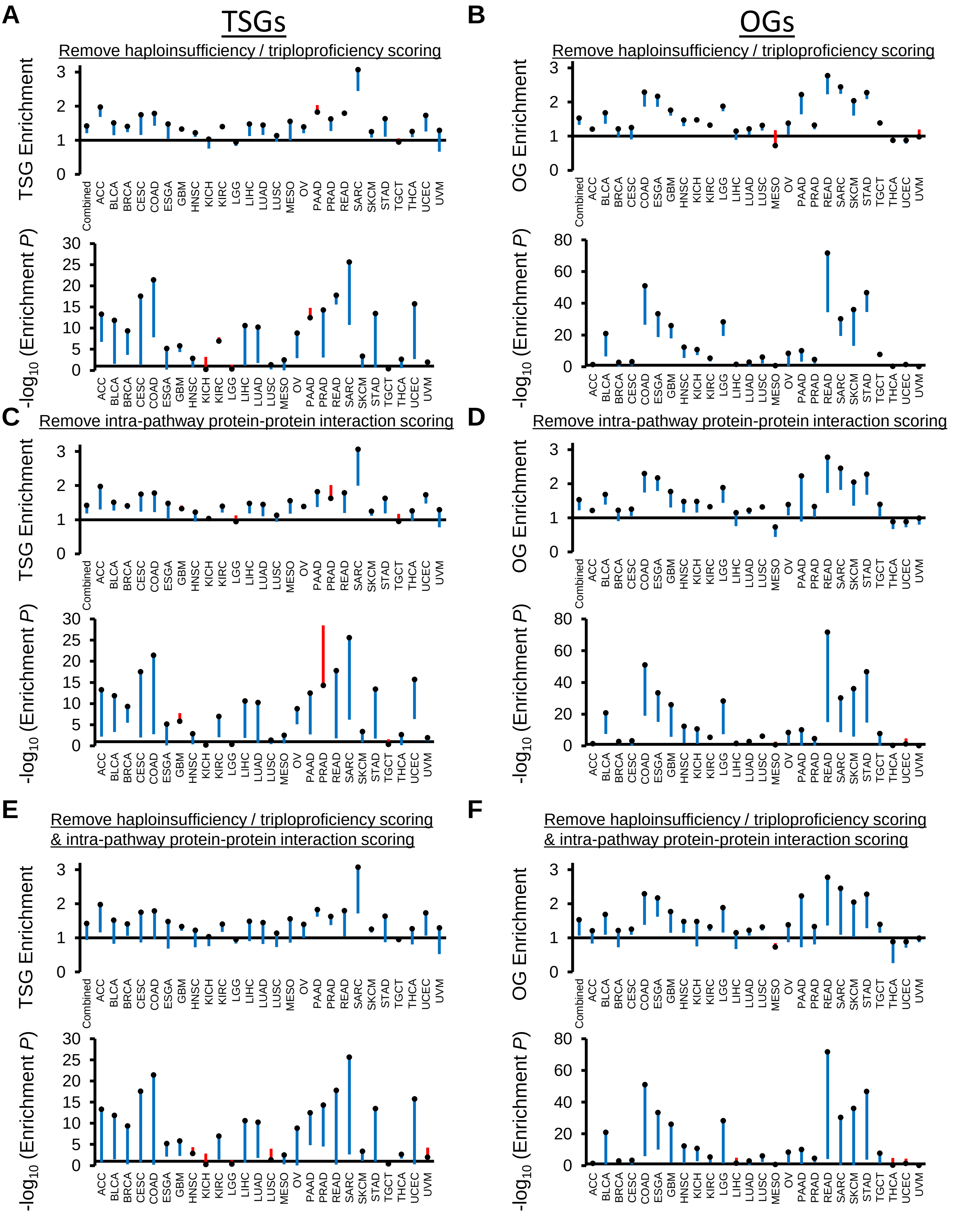

**Supplementary Figure 3. Contribution of Phenotypes to SWAN Analysis.** (**A-F**) The SWAN default result is plotted as black squares for each tumor type. Changes in tumor suppressor gene (TSG) or oncogene (OG) enrichment are indicated by loss of enrichment (blue) or gain in enrichment (red). See Methods for description of enrichment values and statistics.

### Supplemental Figure 4:
### Example of SWAN analysis of standard molecular biology datasets

**A**

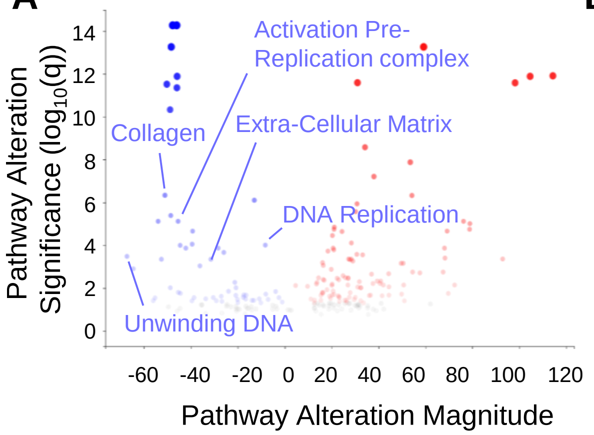


**B**

| Reference | BET Inhibition Strategy | Cell Line | SWAN *P*-Value for Reactome: Activation of the Pre-Replication Complex |
|---|---|---|---|
| Suzuki *et al*. Mol. Med. Rep. (2020) | JQ1 | Human primary myofibroblasts | $5.79 \times 10^{-6}$ |
| Garcia-Carpizo *et al*. Epigenetics Chromatin (2018) | JQ1 | K-562 (Human CML) | $1.48 \times 10^{-7}$ |
| Ren *et al*. PNAS (2018) | JQ1 | MDA-MB-231 (Human TNBC) | $9.78 \times 10^{-4}$ |
| Ren *et al*. PNAS (2018) | MS645 | MDA-MB-231 (Human TNBC) | $2.93 \times 10^{-7}$ |
| Nagaraja *et al*. Nucleic Acids Res (2017) | JQ1 | MCF10A (Human mammary gland) | $1.81 \times 10^{-9}$ |
| Nagaraja *et al*. Nucleic Acids Res (2017) | siBRD4 | MCF10A (Human mammary gland) | $5.10 \times 10^{-8}$ |

**C**

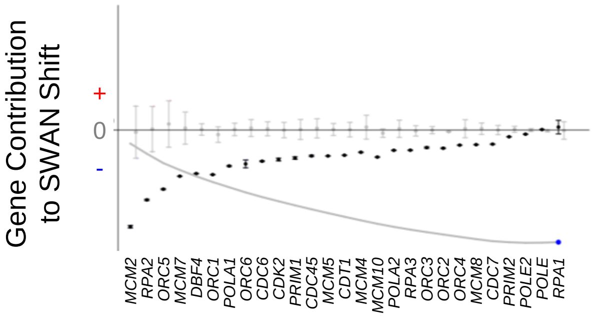


**Supplemental Figure 4: Example of SWAN analysis of standard molecular biology datasets.** BET inhibition reduces transcription of proteins involved in replication initiation. (**A**) SWAN pan-pathway analysis of RNA-seq data from DMSO or JQ1-treated primary fibroblasts. (**B**) SWAN analyses of the Reactome: Activation of the Pre-Replication Complex pathway. Standard SWAN Wilcoxon ranksum *P*-values are presented. (**C**) SWAN analysis of the Reactome: Activation of the Pre-Replication Complex pathway for scrambled siRNA control or siBRD4-treated MCF10A cells.

### Supplemental Figure 5: SWAN integration of TCGA RNA data

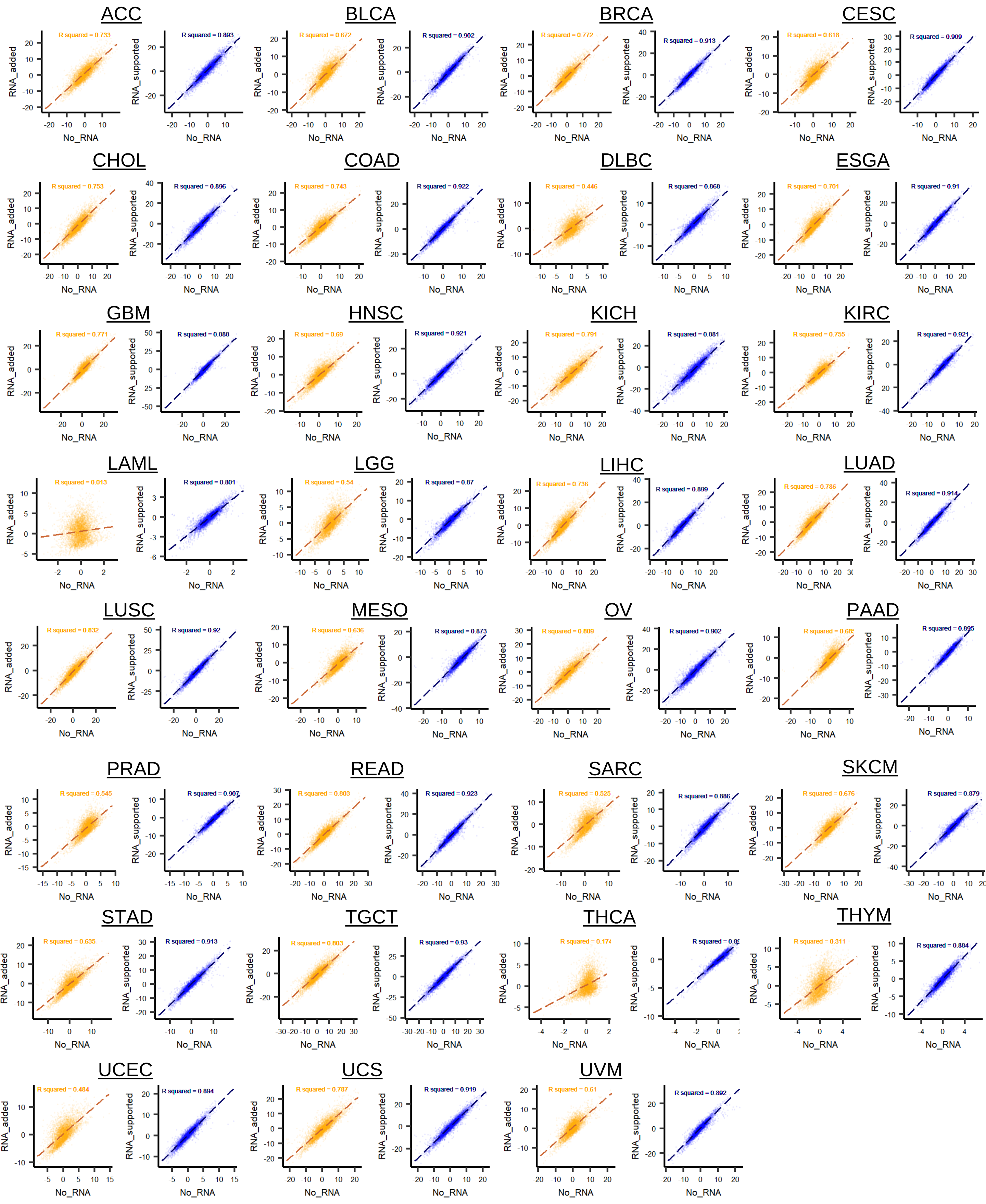

**Supplementary Figure 5. SWAN integration of TCGA RNA data.** SWAN CNA data was integrated with TCGA RNA data using two separate methods. Pan-cancer normalized RNA was added to all CNA data (gold) or added only in cases wherein the CNA and RNA change was detected in the same positive or negative direction (blue). SWAN shifts are plotted without the RNA data (x-axis) and compared to the SWAN shifts with the integrated RNA data (y-axis). An R-squared value is shown for the linear trendline.

### Supplemental Figure 6: Novel oncogenic alterations

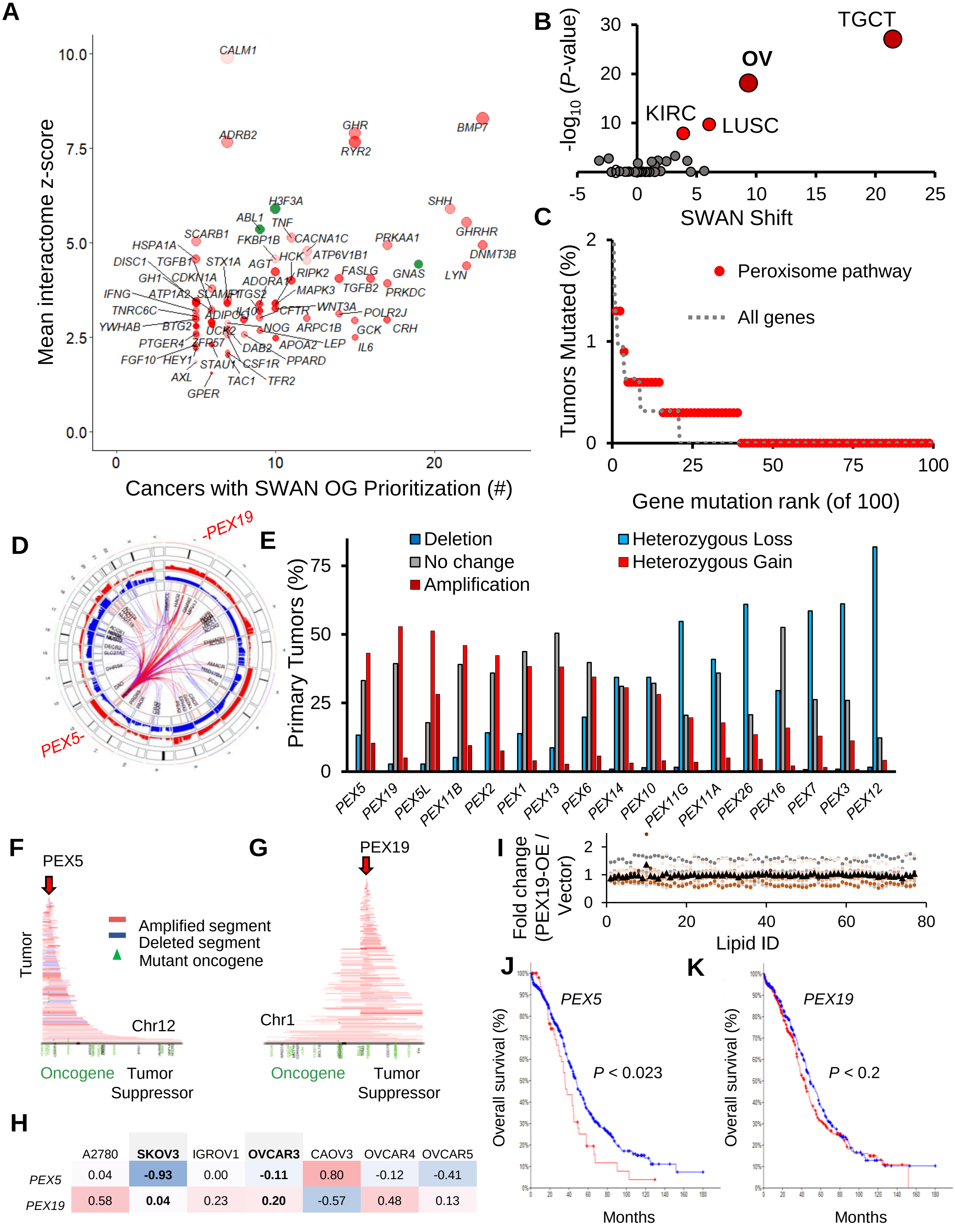


H

| | A2780 | **SKOV3** | IGROV1 | **OVCAR3** | CAOV3 | OVCAR4 | OVCAR5 |
|---|---|---|---|---|---|---|---|
| *PEX5* | 0.04 | **-0.93** | 0.00 | **-0.11** | 0.80 | -0.12 | -0.41 |
| *PEX19* | 0.58 | **0.04** | 0.23 | **0.20** | -0.57 | 0.48 | 0.13 |

**Supplementary Figure 6. Novel oncogenic alterations.** (**A**) Accompanies **Figure 3C**. Potential novel (red) CNA-driven oncogenes along with known COSMIC oncogenes (green). The y-axis refers to SWAN pan-GO-pathway interactome z-scores relative to the number of cancers which included the plotted oncogene in significantly elevated pathways (x-axis). (**B**) SWAN significance of KEGG peroxisome CNA upregulation in the pan-cancer dataset compared to the SWAN shift. (**C**) Mutation rates of all genes in OV are compared to mutation rates of genes within the peroxisome pathway, indicating peroxisome genes are not elevated in SNV mutation rates. (**D**) SWAN Circos plot of the peroxisome pathway in OV, highlighting location of *PEX5* on Chr12p and *PEX19* on Chr1q. (**E**) CNA frequencies of all genes within the KEGG peroxisome pathway in OV. (**F**) CAIRN plot of CNAs overlapping *PEX5* in OV. (**G**) CAIRN plot of CNAs overlapping *PEX19* in OV. (**H**) $Log_2$ CNAs of CCLE-analyzed ovarian cancer cell lines for tested peroxisome genes, highlighting the rationale for usage of SKOV3 and OVCAR3 for molecular investigation. (**I**) Unbiased ultra-performance liquid chromatography mass-spectrometry metabolomics of analyzed lipids (see Methods, **Table S4**) in SKOV3 cells. Orange circles are individual replicates of PEX19-OE relative to the mean control cells. Grey circles are individual replicates of control cells relative to the mean of control cells. Black triangles are means of PEX19-OE fold changes. No changes were nominally significant, $P < 0.05$ by t-test. (**J**) Overall survival of *PEX5* upregulated tumors, by KmPlot of ovarian cancer data. (**K**) Overall survival of *PEX19* upregulated tumors, by KmPlot of ovarian cancer data.

### Supplemental Figure 7: Differential CNA pathways and survival

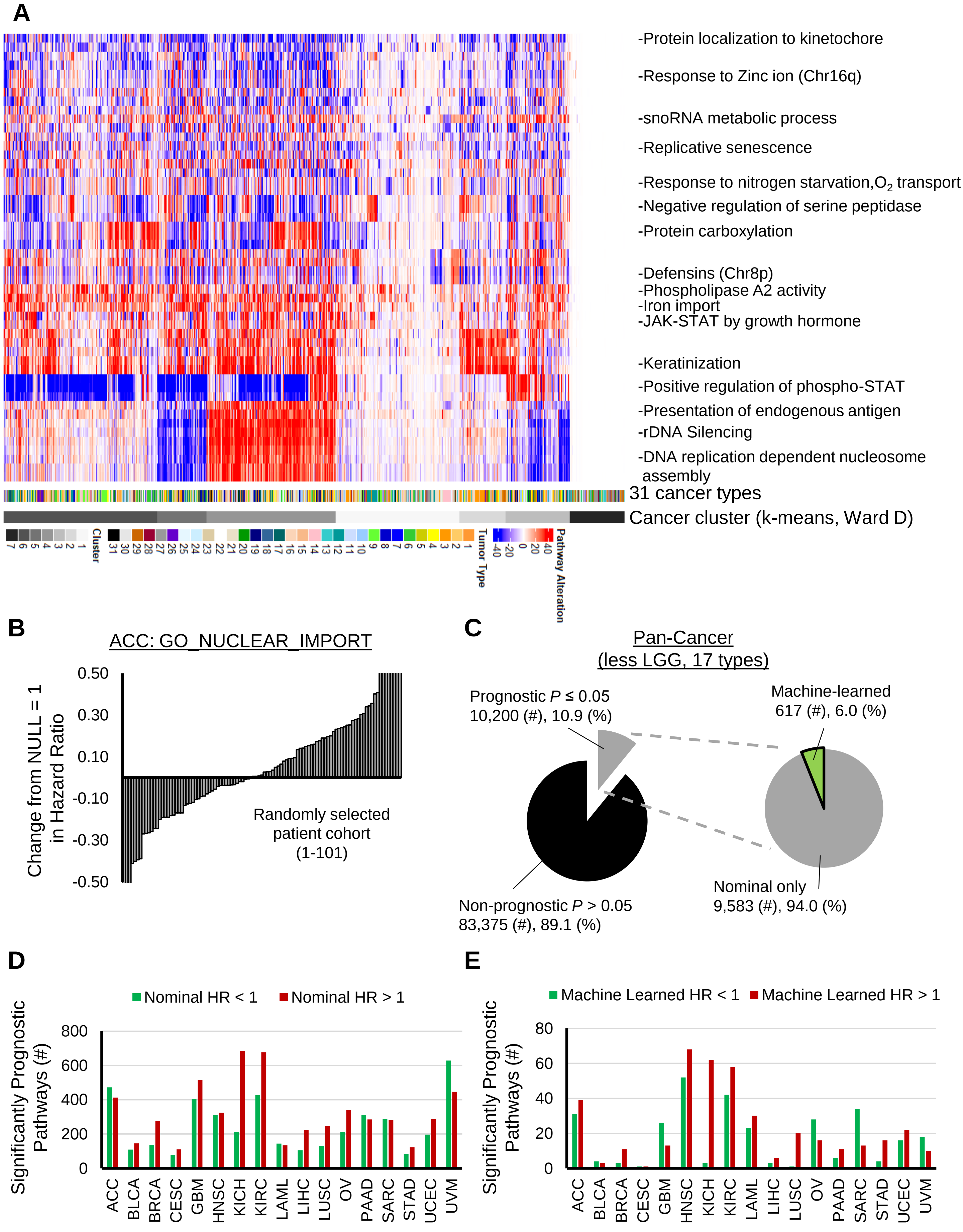

**Supplementary Figure 7. Differential CNA pathways and survival.** (**A**) K-means clustering of the top 1% differentially SWAN shifted pathways. Per-tumor SWAN shifts are displayed for all tumors analyzed. (**B**) Cox-proportional hazard ratios calculated from 101 randomly picked subsets of ACC patients with high or low SWAN shifts within the GO Nuclear Import pathway (see Methods). As shown, the subset of patients analyzed strongly affects the interpretation of survival prognosis. Y-axis was capped at ±0.5 to enable visualization of mid-range random subsets. (**C**) Pie charts showing the proportion of pathways reported as nominally prognostic as the full patient cohort (left) compared to those which are provided as confidently machine-learned prognostic (right). LGG was graphically omitted here since its consistent CNA patterns are an outlier, which contributes an unusually large portion of prognostic pathways. (**D**) Summary of $P \leq 0.05$ prognostic pathways per compatibly analyzed tumor type (LGG omitted). (**E**) Summary of prognostic pathways per compatibly analyzed tumor type (LGG omitted) after machine learning process was performed.
